# MYC disrupts transcriptional and metabolic circadian oscillations in cancer and promotes enhanced biosynthesis

**DOI:** 10.1101/2023.01.03.522637

**Authors:** Juliana Cazarin, Rachel E. DeRollo, Siti Noor Ain Binti Ahmad Shahidan, Jamison B. Burchett, Daniel Mwangi, Saikumari Krishnaiah, Annie L. Hsieh, Zandra E. Walton, Rebekah Brooks, Stephano S. Mello, Aalim M. Weljie, Chi V. Dang, Brian J. Altman

## Abstract

The molecular circadian clock, which controls rhythmic 24-hour oscillation of genes, proteins, and metabolites in healthy tissues, is disrupted across many human cancers. Deregulated expression of the MYC oncoprotein has been shown to alter expression of molecular clock genes, leading to a disruption of molecular clock oscillation across cancer types. It remains unclear what benefit cancer cells gain from suppressing clock oscillation, and how this loss of molecular clock oscillation impacts global gene expression and metabolism in cancer. We hypothesized that MYC or its paralog N-MYC (collectively termed MYC herein) suppress oscillation of gene expression and metabolism to upregulate pathways involved in biosynthesis in a static, non-oscillatory fashion. To test this, cells from distinct cancer types with inducible MYC were examined, using time-series RNA-sequencing and metabolomics, to determine the extent to which MYC activation disrupts global oscillation of genes, gene expression pathways, and metabolites. We focused our analyses on genes, pathways, and metabolites that changed in common across multiple cancer cell line models. We report here that MYC disrupted over 85% of oscillating genes, while instead promoting enhanced ribosomal and mitochondrial biogenesis and suppressed cell attachment pathways. Notably, when MYC is activated, biosynthetic programs that were formerly circadian flipped to being upregulated in an oscillation-free manner. Further, activation of MYC ablates the oscillation of nutrient transporter proteins while greatly upregulating transporter expression, cell surface localization, and intracellular amino acid pools. Finally, we report that MYC disrupts metabolite oscillations and the temporal segregation of amino acid metabolism from nucleotide metabolism. Our results demonstrate that MYC disruption of the molecular circadian clock releases metabolic and biosynthetic processes from circadian control, which may provide a distinct advantage to cancer cells.

## Introduction

Circadian rhythms are ∼24-hour rhythms that occur in many organisms and can be observed on the cellular level to rhythmically control transcription, protein levels, and metabolic processes. Recent analyses have revealed that many human cancers have disrupted or ablated circadian clocks. However, what impact circadian disruption has on cancer cell transcriptional and metabolic programming has not been determined.

Circadian rhythms entrain organisms’ activity (such as sleep-wake cycles) and metabolism (such as segmentation of catabolism and biosynthesis) to the day / night cycle. The ‘central clock’, housed within the suprachiasmatic nucleus of the hypothalamus, receives light signals through the eye and communicates these signals to peripheral cell-autonomous ‘molecular clocks’ present in every cell and tissue, which control oscillatory gene expression [1]. The molecular clock is controlled by the basic helix-loop-helix (bHLH) transcription factors CLOCK and BMAL1 (encoded by *ARNTL*), which, as a heterodimer, rhythmically regulate target gene expression in an oscillatory fashion [1]. CLOCK and BMAL1 control their own activity and levels through a series of feedback loops. In the first and most central loop, rhythmically produced PER and CRY proteins inhibit the activity of CLOCK and BMAL1, leading to antiphase oscillation of CLOCK-BMAL1 activity and PER and CRY levels and localization. In a second accessory loop required for molecular clock function in many cells and tissues, the nuclear hormone receptors and transcriptional repressors REV-ERBα and REV-ERBβ compete with the transcriptional ROR activators to control rhythmic transcription of BMAL1 (*ARNTL*) ([1-3]). The molecular clock is thus responsible for the rhythmic regulation of thousands of transcripts in mouse and primate, comprising more than 50% of protein coding transcripts in some tissues [4, 5]. Oscillatory gene expression arises from direct action of CLOCK-BMAL1 on promoters and enhancers, from binding of the REV-ERB and ROR family to promoters, and from action of secondary tissue-specific transcription factors that themselves are rhythmically controlled by the molecular clock [6, 7]. Importantly, not all oscillation in a cell arises solely from mRNA: several studies have shown that proteins and metabolites also oscillate in a manner not directly dependent on mRNA oscillation [8-10].

The molecular circadian clock is tumor suppressive in many tissues. Indeed, disruption of the molecular clock, either by behavioral disruption or genetic mutation, can accelerate or initiate tumorigenesis in lung, liver, bone, colon, pancreas, melanocyte, and other tissue types [11-20]. Supporting the notion that the clock is tumor suppressive in human cancer, circadian gene expression is often disrupted in tumor tissue as compared to normal tissue [21]. Perhaps more importantly, three independent computational analyses each revealed that normal progression and oscillation of the circadian clock is dampened or lost in human tumor tissue compared to normal tissue [22-24]. We and others have proposed that clock disruption in cancer may release cellular processes such as cell cycle or metabolic fluxes from circadian control to be constantly and statically up- or downregulated [25, 26]. Supporting this notion, breast cancer patients with mutations in molecular clock genes have a worse survival rate than those with wild-type alleles [21]. However, given that mutations in individual molecular clock genes are rare [21], it remains unclear how circadian disruption arises in cancer cells, and what formerly circadian programs lose oscillation as a result of this disruption.

A well-defined source of circadian disruption in cancer is amplification of the *MYC* oncogene or its closely related paralogue *MYCN* (N-MYC), which leads to overexpression of MYC or N-MYC. Amplification of at least one *MYC* family member is quite common in human cancer, overall occurring in nearly one third of all cases [27]. The MYC-family proteins are E-box binding transcription factors which, when amplified, tend to drive continued transit through the cell cycle, and to upregulate nutrient uptake, protein translation, and biomass accumulation, amongst many other functions [27-29]. Amplified MYC has also been proposed to increase overall transcriptional output [29, 30]. Amplified MYC and N-MYC have been shown in multiple cancer models to dampen or ablate molecular clock gene oscillation [31-36]. We identified a mechanism whereby MYC and N-MYC directly upregulate the REV-ERB proteins, leading to BMAL1 suppression and a collapse of molecular clock gene oscillation [31, 32]. We also observed that cell-autonomous oscillation of glucose was disrupted by MYC [32]; however, it remained unclear which global transcriptional and metabolic programs that are normally circadian controlled were disrupted by MYC amplification. Since MYC drives enhanced nutrient uptake and biosynthesis across multiple cancer, we hypothesized that MYC disruption of the molecular clock releases these processes from circadian control to instead be enhanced by amplified MYC.

In this study, we utilize multiple cell line models representing neuroblastoma and osteosarcoma to determine the role of MYC overexpression in disruption of circadian oscillations of transcription and cell-autonomous metabolism. We perform time-series analysis to identify which transcriptional programs are circadian in the absence of MYC across multiple cell lines; and to also determine, in a time-independent fashion, which genes and pathways MYC upregulates or suppresses in common between the three cell models. We next combine these two analyses to determine which gene expression programs and pathways switch from being oscillatory to being up- or downregulated when MYC is activated. In our analysis, we have taken an agnostic approach, presenting the most significantly enriched genes and pathways rather than choosing specific pathways of interest. Using this agnostic approach, we determine that across multiple cell lines, pathways associated with metabolism and biosynthesis lose oscillation and become upregulated by MYC in a static, oscillation-independent fashion. Finally, we examine which metabolic circadian cycles are disrupted by MYC, and how these connect to changes in the oscillation and expression of nutrient transporters.

## Results

### Oncogenic MYC ablates global transcriptional oscillation

We and others have shown that ectopic MYC disrupts the molecular clock machinery across several types of cancer models [31-36]. Since many of the components of the molecular clock pathway, including CLOCK, BMAL1, and the REV-ERB proteins are transcription factors, it can be surmised that disruption in normal oscillation of the molecular clock may lead to loss of rhythmicity of global circadian output. Similarly, MYC may upregulate genes and programs formerly regulated by the molecular clock. Therefore, the global transcriptional impact of loss of molecular clock gene expression rhythmicity remained unclear. To address this, we utilized three separate cancer cell lines we previously characterized to have intact molecular clocks that are disrupted by overexpressed MYC or N-MYC: SHEP N-MYC-ER (neuroblastoma), SKNAS N-MYC-ER (neuroblastoma), and U2OS MYC-ER (osteosarcoma) [31, 32]. All three lines have inducible overexpressed MYC-ER or N-MYC-ER that is activated by 4-hydroxytamoxifen (4OHT). In this model, MYC-ER or N-MYC-ER is constitutively expressed, and when exposed to 4OHT, translocates to the nucleus to lead to upregulation of target genes [31, 37]. Activated MYC-ER and N-MYC-ER will henceforth be referred to for all cell lines as MYC-ON. When overexpressed MYC-ER is inactive in control conditions, this condition is referred to as MYC-OFF; these cells are treated with ethanol as a vehicle control. We quantified the degree of MYC or N-MYC overexpression in all 3 cell lines, by qPCR (SHEP and SKNAS) and by immunoblot (U2OS). We found that N-MYC-ER was overexpressed ∼750-fold in SHEP and ∼7000 fold in SKNAS compared to endogenous N-MYC in SKNAS (SHEP do not express N-MYC) (**Supplemental Figure 1A**), while MYC-ER was overexpressed ∼5.5-fold in U2OS, (**Supplemental Figure 1B**). We performed time-series qPCR analysis on these three cell lines with or without MYC activation in n=2 biological replicates per cell line: cells were entrained with dexamethasone at CT0, and collected from CT24-72 or 76 at 2-4 hour intervals. CT refers to ‘circadian time’, the number of hours after dexamethasone entrainment. qPCR analysis revealed that in all three lines, MYC elevated *PER2* and *NR1D1* (REV-ERBα) while suppressing *ARNTL* (BMAL1) (**Figure 1**). In addition, we used the ECHO algorithm [38] to detect oscillation in our time-series qPCR. MYC and N-MYC ablated rhythmicity of REV-ERBα in SHEP and U2OS, BMAL1 in SHEP and SKNAS, and *PER2* in SKNAS. This agrees with prior findings from our lab and others that MYC and N-MYC dampen or ablate oscillation of core molecular circadian clock genes [31-36]. In contrast, we previously showed that tamoxifen on its own does not affect the molecular clock, nor does expression of a MYC-ER protein that lacks transcriptional activity [31].

**Figure 1.**
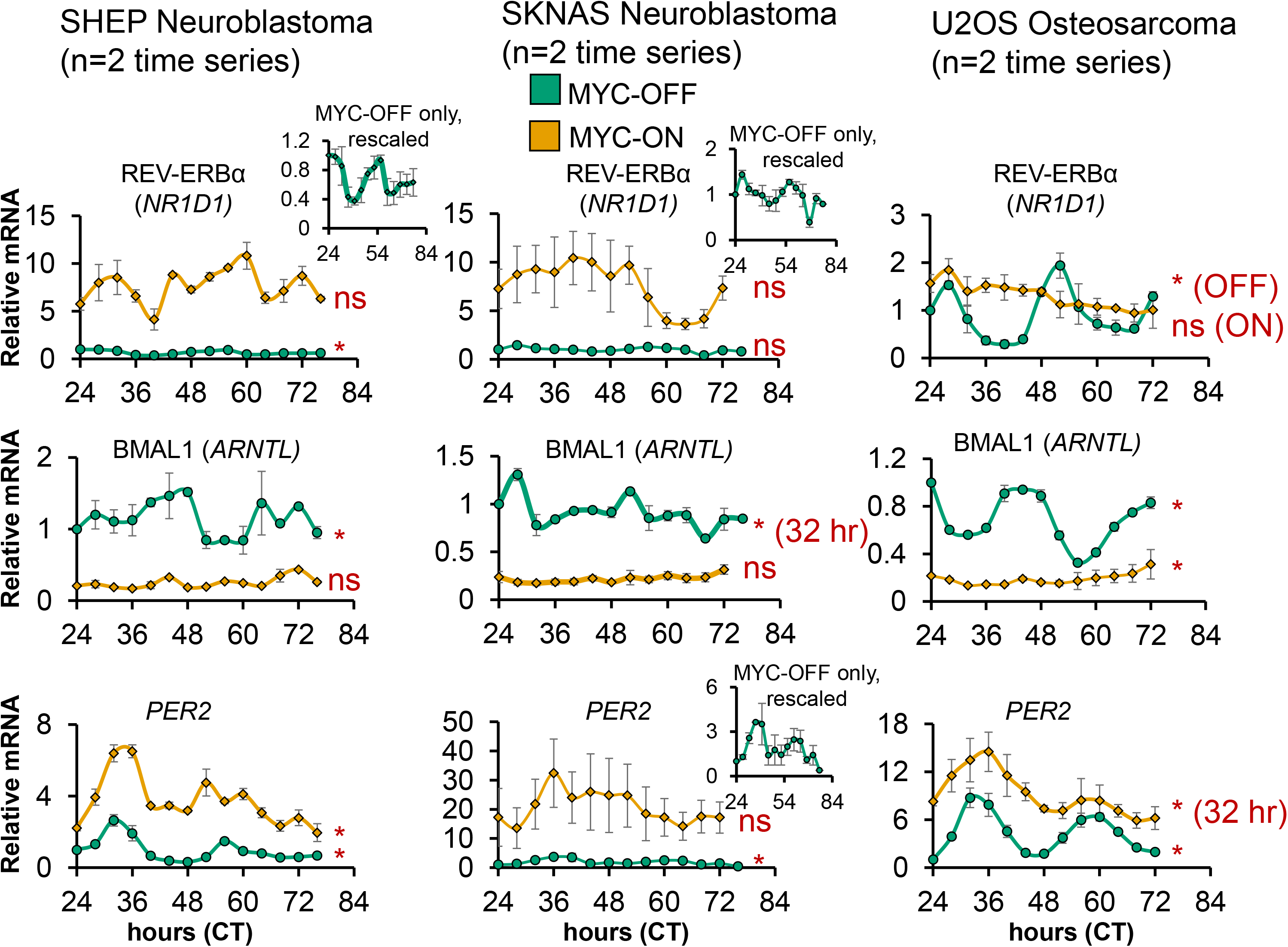
Oncogenic MYC disrupts molecular clock gene oscillation in cancer cells. SHEP N-MYC-ER and U2OS MYC-ER were treated with ethanol control (MYC-OFF) or 4-hydroxytamoxifen (MYC-ON) (4OHT) to activate MYC, and entrained with dexamethasone, and after 24 hours, RNA was collected every 2-4 hours for the indicated time period. Expression of the indicated genes was determined by quantitative PCR (qPCR), normalized to β2M. n=2 biological replicates were performed, with error bars representing standard error of the mean (S.E.M.). CT = circadian time. Note the inset MYC-OFF only graphs for SHEP *NR1D1*, and SKNAS *PER2,* to show oscillation of these genes in MYC-OFF cells on a different scale. qPCR results were analyzed for rhythmicity by ECHO for both MYC-OFF and MYC-ON, with genes with a p value of < 0.05 and a BH.Adj.P.Value < 0.05 deemed rhythmic. For those with a period outside the 20-28 hour range, this is noted.

Our MYC-ER and N-MYC-ER cell lines model acute MYC overexpression, but do not show whether inhibiting MYC in amplified settings also affects clock gene expression. We previously showed in models of MYC amplification representing liver cancer (tumors and cell lines) and Burkitt’s Lymphoma that suppressing amplified MYC leads to changes in clock gene expression, including increasing BMAL1 (which suggested that MYC suppressed BMAL1 in these settings) [31, 32]. To further query the how suppressing elevated MYC in cancer cells may affect circadian gene expression, we leveraged publicly available data where PC3 prostate cancer cells, known to have high endogenous (non-amplified) MYC levels [39, 40], were treated with the new generation MYC inhibitor MYCi361 [41]. Using differential expression analysis with DeSeq2 [42], we found that in cells treated with MYCi361, BMAL1 was upregulated and *PER2* was downregulated, suggesting that MYC suppresses BMAL1 and upregulates *PER2* in these cells, similar to in SHEP, SKNAS, and U2OS (**Supplemental Figure 1C**). In contrast, MYCi361 led to upregulation of *NR1D1* (REV-ERBα) and *NR1D2* (REV-ERBβ) suggesting that in these cells, MYC suppressed the REV-ERBs (**Supplemental Figure 1C**). While the mechanism for how *MYC* affects expression of molecular clock genes in these cells remains unclear, these findings show that suppression of naturally elevated endogenous MYC changes molecular clock gene expression.

We next performed RNA-sequencing of each circadian time series in the three cancer cell lines with or without MYC activation, as replicates, and used the ECHO algorithm [38] to detect oscillating genes. Depending on cell line, we detected between ∼700-1600 oscillating genes in the MYC-OFF condition (**Figure 2A**). To determine how the strength of these oscillations in MYC-OFF compared to established primary cell models of circadian oscillation, we downloaded publicly available time-series datasets of entrained primary macrophages (Mϕ) and mouse embryonic fibroblasts (MEFs) in constant, free-running conditions after entrainment [9, 43], and analyzed them with ECHO. For each analysis, we compared median amplitude of all oscillating genes. We found that the median amplitude of oscillating genes in MYC-OFF in U2OS, SHEP and SKNAS was between ∼0.5 and 1.7, which was within the range of what was observed in primary macrophages (0.35) and MEFs (1.7) (**Supplemental Figure 2A**), suggesting that the transcriptional oscillations we observed in cancer cell lines were of a similar amplitude to those in primary cells. We next asked which specific genes oscillated in common in MYC-OFF SHEP, SKNAS, and U2OS cells. Supporting the notion that the identity of circadian output genes is highly variable between different cell and organ systems [4, 5], there was little overlap in the specific identity of oscillating genes in the MYC-OFF conditions amongst the three cell lines, with only 3 molecular clock genes, *CRY2, PER2, and PER3*, oscillating in common in all 3 cell lines. (**Figure 2B**). Nonetheless, when we examined these oscillatory genes in the MYC-ON condition, we found that 90% or more of genes that oscillated in MYC-OFF no longer oscillated in MYC-ON (**Figure 2A**). This suggested a global collapse of the normal transcriptional oscillatory program in the presence of ectopic MYC.

**Figure 2.**
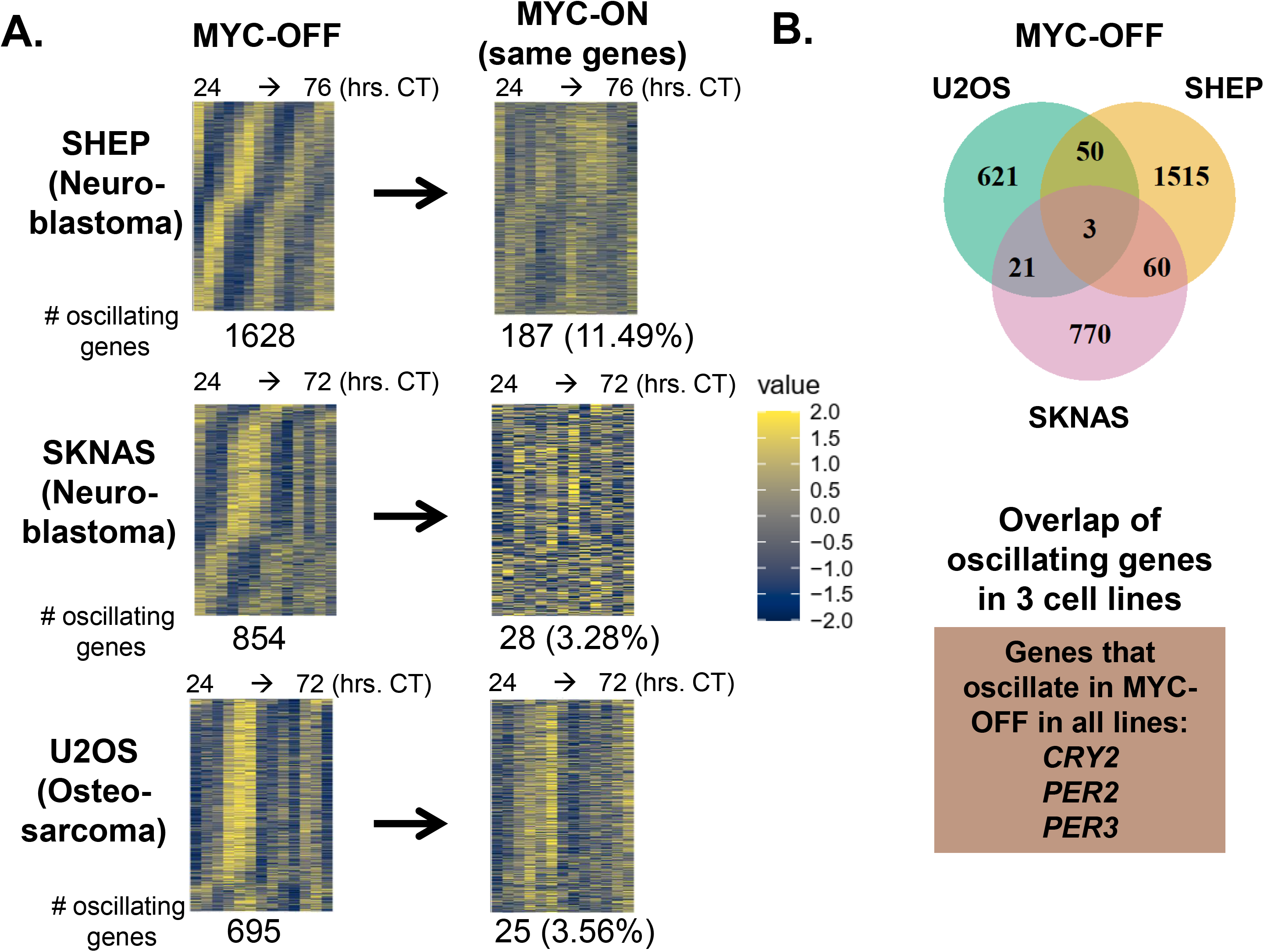
Oncogenic MYC disrupts global transcriptomic circadian oscillation in cancer cells. **A.** RNA-sequencing was performed on the same n=2 replicate SHEP N-MYC-ER, SKNAS-N-MYC-ER or U2OS MYC-ER time series ± 4OHT and + dexamethasone shown in Figure 1, with RNA samples collected every 2-4 hours at the indicated timepoints. RNA was analyzed for rhythmicity by ECHO for both MYC-OFF and MYC-ON, with genes with a 20-28 hour period and with p value < 0.05 and a BH.Adj.P.Value < 0.05 (SHEP, U2OS) or 0.1 (SKNAS) deemed rhythmic. These genes were sorted by phase and are presented in a heatmap for MYC-OFF. For MYC-ON, the same genes that are rhythmic in MYC-OFF are presented in the same order, but with MYC-ON values instead. **B.** The overlap of oscillating genes in MYC-OFF identified in **(A)** is shown by Venn diagram. Those genes that oscillated in all 3 cell lines are highlighted below the Venn diagram.

Surprisingly, when MYC was activated, many genes gained oscillation that were previously not oscillatory. We found that MYC-ON cells contained between ∼500-1500 oscillating genes, with approximately 90% not oscillating in MYC-OFF (**Supplemental Figure 2B**). MYC suppresses BMAL1 expression (**Figure 1** and [31-34]), and it was previously suggested that oscillations occurring in low or absent BMAL1 may be of very low amplitude [44, 45]. To test whether this was occurring in our cell line models, we examined the amplitude of all genes oscillating with a circadian period in MYC-OFF and MYC-ON (**Supplemental Figure 2C**). We found no particular tendency of oscillating genes to be of lower amplitude in MYC-ON across all three cell lines. Overall, these results showed that activation of ectopic MYC suppressed normal circadian oscillatory output across multiple independent models.

### Oncogenic MYC causes a shift in the number and identity of oscillating transcriptional programs

While the individual identity of oscillating genes varies widely between tissues, there are basic cellular functions that are nonetheless commonly circadian-regulated across multiple tissues and cell types [4, 5]. For instance, it was shown in baboon that metabolic, vesicle-trafficking, and cell membrane and junction gene expression programs are commonly rhythmic in many different tissues [5]. Given the lack of overlap in the specific identity of oscillating genes in our three cell line models in MYC-OFF conditions, we next asked which transcriptional programs oscillated in each of these cells. To accomplish this, we used complementary methods of phase-dependent and phase-independent analysis. Phase-dependent analysis assumes that mRNAs in a given pathway or group peak together in the same circadian phase, and thus groups genes that peak together for pathway analysis. We used the phase-dependent algorithm Phase Set Enrichment Analysis (PSEA) to determine which gene sets and pathways were enriched in oscillating genes in MYC-OFF and MYC-ON in a phase-dependent fashion [46]. To determine when these programs peaked, we plotted significantly enriched oscillating programs on radial histograms, which circularize the X-axis to accurately show a repeating circadian time scale of CTs. Similar to individual genes in **Figure 2**, most of the enriched oscillatory programs in MYC-OFF did not oscillate in MYC-ON (**Figure 3A**). Intriguingly, even when oscillation was maintained from MYC-OFF to MYC-ON, the time in which these programs peaked was altered. For instance, in SHEP MYC-OFF, most gene expression programs detected peaked between CT 17-21, while in MYC-ON, those that were shared instead peaked mostly between CT 6-15. In SKNAS, only a single gene set amongst the 351 sets detected by PSEA to rhythmically peak in MYC-OFF continued to oscillate in MYC-ON. U2OS osteosarcoma were similar to SHEP: in MYC-OFF, the strongest peak occurred at CT16, while no shared gene sets peaked at this time in MYC-ON. The gene sets determined as being oscillatory in MYC-ON also had different timing of peak expression compared to MYC-OFF (**Supplemental Figure 3A, compare to Figure 3A**). We focused on which gene expression programs were shared by at least two cell lines and ranked these according to average significance score across the cell lines. Similar to previous findings in baboon, in MYC-OFF cells, those shared between cell lines corresponded to endoplasmic reticulum and cell junction and contact, as well as cell cycle / DNA replication (**Figure 3B**). In contrast, there was far less overlap of oscillating gene expression programs in MYC-ON (**Supplemental Figure 3B**), and no identifiable common themes emerged. This suggested that some generalities exist in SHEP, SKNAS, and U2OS without elevated MYC; while by contrast, MYC overexpression does not drive a common oscillatory program at the transcript level in different cell lines.

**Figure 3.**
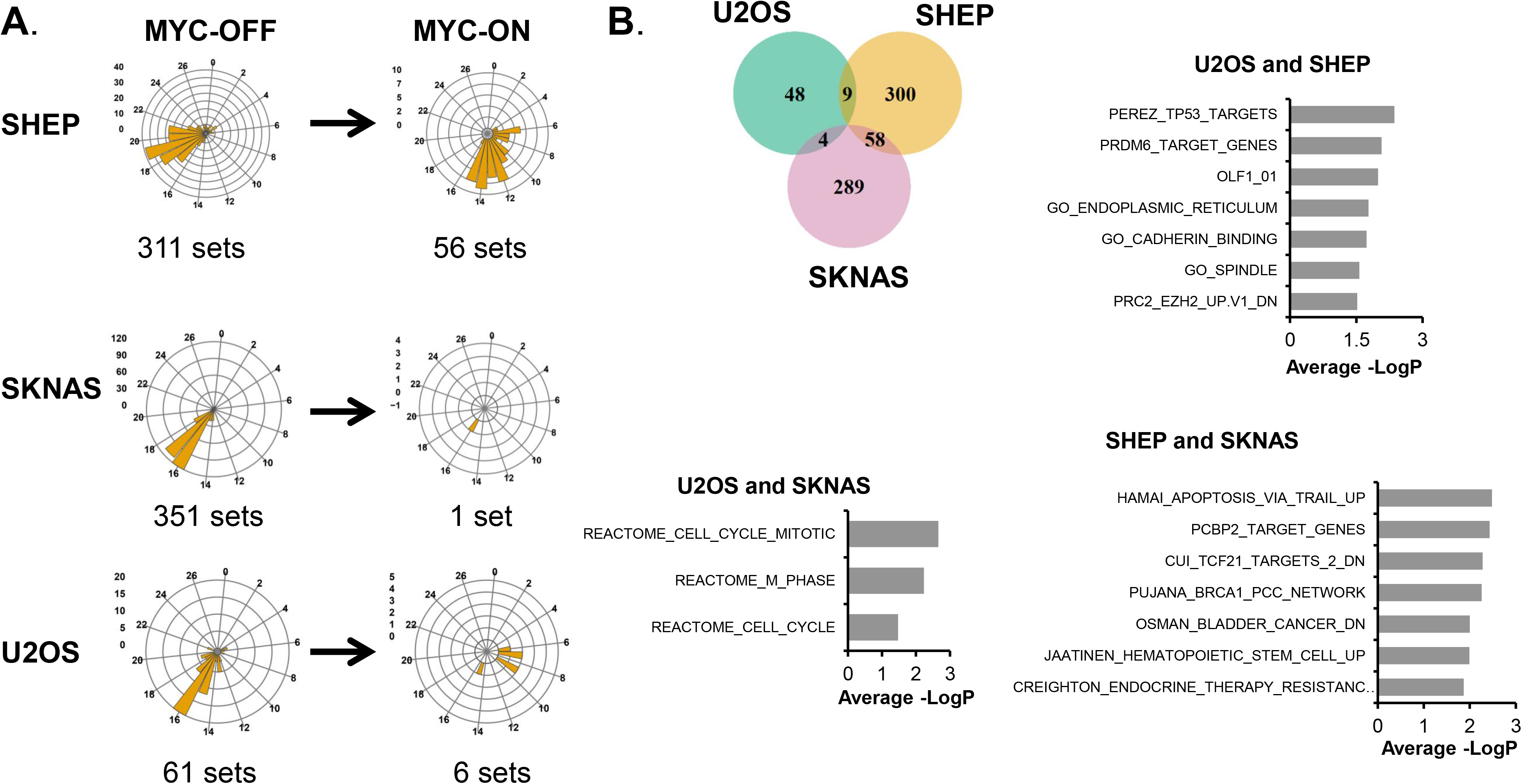
Oncogenic MYC shifts the identify of oscillating transcriptional programs. **A.** Oscillatory genes in SHEP N-MYC-ER, SKNAS-N-MYC-ER or U2OS MYC-ER were analyzed with Phase Set Enrichment Analysis (PSEA) for oscillatory pathway enrichment, with a period of 20 – 28 hours and q-value (vs. background) < 0.2 and p-value (vs. background) <0.1 deemed significant. Oscillatory pathways were binned by hour and plotted on a polar histogram, which is a circular histogram that accurately displays a repeating circadian scale on the X-axis. The Y-axis scale for each histogram is on the left side. MYC-OFF significant pathways are shown, and the same pathways are also plotted for MYC-ON. **B.** Overlap of oscillatory programs in MYC-OFF are shown by Venn diagram, and identity of oscillatory programs are plotted by average of –LogP from each cell line of overlap. The most highly significant pathways (up to 10) for each overlap are shown.

We also utilized a phase-independent analysis methodology to determine which transcriptional programs were enriched amongst oscillatory genes. Some oscillatory programs, including cell cycle and the molecular circadian clock itself, have genes that may peak at different times of the day [47, 48], making detection of these programs with PSEA less efficient. Thus, we analyzed oscillatory genes in each cell line regardless of their peak of oscillation for pathway enrichment with ToppFun functional enrichment suite, which combines several different enrichment libraries [49]. We found that, despite the fact that there were largely similar numbers of oscillating genes between MYC-OFF and MYC-ON, there tended to be less enriched gene sets among MYC-ON. In SHEP MYC-OFF, oscillating genes were enriched for endosomal and lysosomal trafficking, and in U2OS and SKNAS, programs related to the molecular circadian clock were enriched (**Table 1**). In contrast, in SHEP and U2OS MYC-ON, programs related to cell cycle were instead enriched, while no programs were significantly enriched in SKNAS MYC-ON (**Table 1**).

We further investigated this apparent gain in cell cycle gene rhythmicity in SHEP and U2OS MYC-ON by interrogating which cell cycle genes were oscillatory with a ∼24-hour period in the absence or presence of activated MYC. In both cell lines, there were fewer significantly rhythmic cell cycle genes in MYC-OFF (11 in SHEP, 3 in U2OS) as compared to MYC-ON (16 in SHEP, 5 in U2OS, **Supplemental Figure 3C,D**), suggesting a potential gain in cell cycle gene oscillation in the presence of MYC as circadian rhythmicity is diminished. This agrees with prior findings in U2OS that intentional disruption of the molecular clock promotes enhanced cell cycle advance [17]. Overall, these data suggested that ectopic MYC reduces or abrogates rhythmicity of gene expression programs that normally are controlled by the molecular clock, and instead may promote cell cycle rhythmicity.

### Oncogenic MYC promotes a distinct transcriptional program of biosynthesis and loss of attachment across multiple cell models

We next sought to address which genes and pathways MYC up- or downregulated, without regards to rhythmicity. MYC can have many disparate functions in cancer, which may depend on 1) the degree to which MYC is overexpressed, 2) which promoters and enhancers it binds, 3) if MYC cooperates with other transcription factors such as MIZ1 to inhibit expression of certain genes, and 4) if MYC strongly regulates pause release ([27, 30, 50-53]. Our MYC-ER models gave us the opportunity to determine the role of MYC activation across three unrelated cell lines, and to determine commonalities in MYC rewiring global transcription. To accomplish this, we performed differential expression analysis, using DeSeq2, on our time-series RNA-sequencing experiments [42]. Differential expression analysis revealed that MYC activation resulted in the upregulation (by a least 10%) between ∼5500 to ∼7700 genes and suppression (by at least 10%) between ∼5600 to ∼8100 genes (**Figure 4A**), in line with previous reports that MYC exerts a strong genome-wide repressive role [54, 55]. Overall, this corresponded to approximately 30-40% of all genes detected by RNA-sequencing. We performed Gene Set Enrichment Analysis (GSEA) for all genes up- or down-regulated by at least 10%, and observed which gene sets were enriched in common between all three cell lines [56-59]. There were 160 gene sets upregulated in common between the three inducible cell lines. Common programs were focused on canonical MYC targets, mitochondrial gene expression and assembly, ribosome biogenesis, tRNA processing, and metabolism (**Figure 4B**). In contrast, 316 gene sets were downregulated in common between all three cell lines. These suppressed gene expression programs corresponded to cell adhesion, extracellular matrix (ECM), collagen, focal adhesion, and HDAC targets (**Figure 4C**). While MYC has not been previously described to negatively regulate these programs on a global transcriptional scale, their downregulation is consistent with previous reports that MYC may promote loss of attachment and an invasive and metastatic phenotype in cancer cells [60-63].

**Figure 4.**
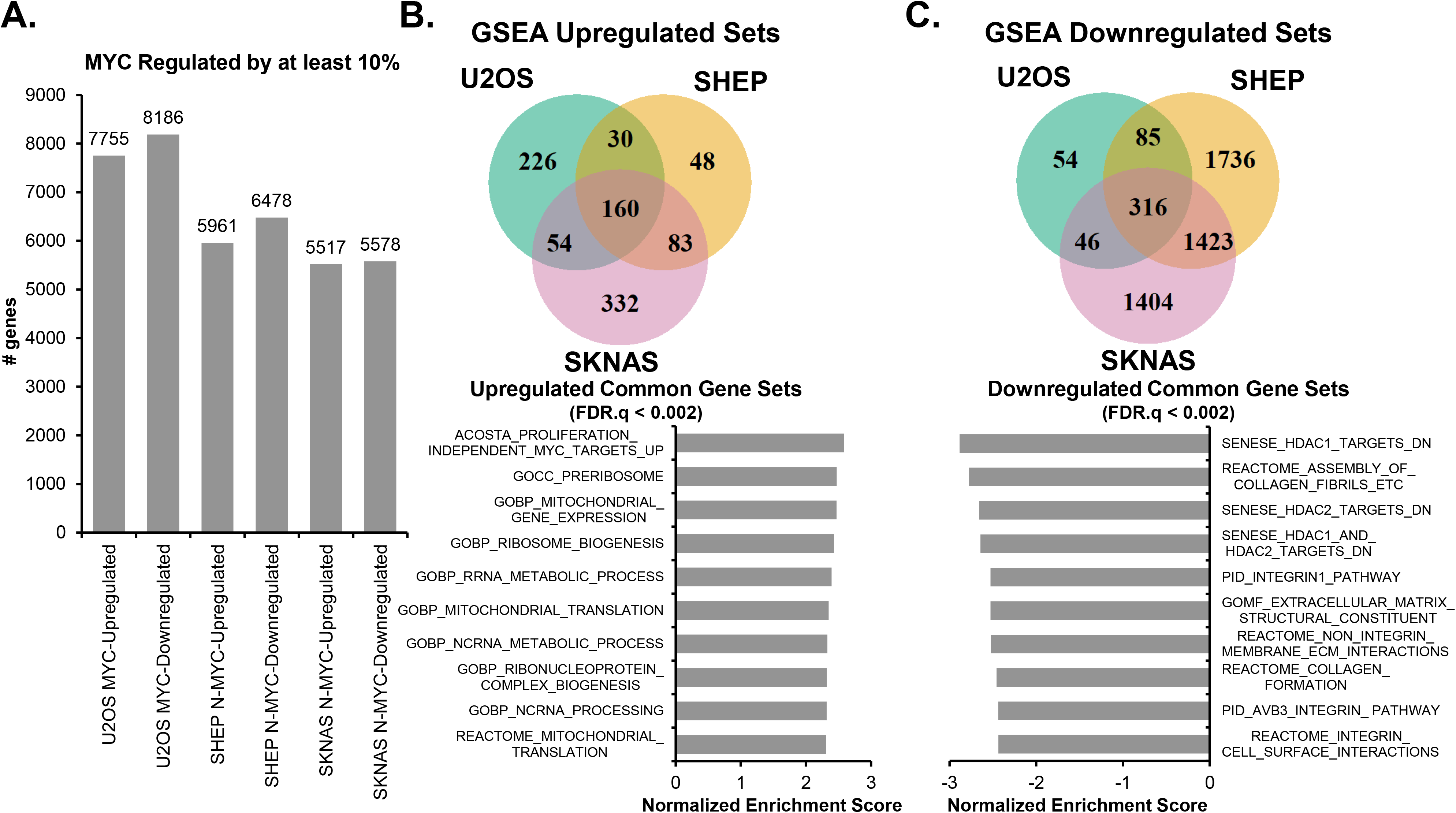
MYC upregulates biosynthetic and metabolic processes and suppresses cell adhesion processes across cancer cell lines. **A.** Differential expression analysis, using DeSeq2, was performed on MYC-ON vs MYC-OFF for SHEP N-MYC-ER, SKNAS-N-MYC-ER or U2OS MYC-ER, with p.adj < 0.05 deemed significant. Genes that were up- or down-regulated in MYC-ON by at least 10% are shown. **B,C.** Gene Set Enrichment Analysis (GSEA) was performed on genes up- or down-regulated by at least 10% in each cell line using GSEA Pre-Ranked, with gene sets with FDR q-val < 0.25 deemed significant. Venn diagrams of overlapping gene sets from MYC-ON upregulated (**B**) and MYC-ON downregulated processes (**C**) are shown, and the most highly significant pathways (up to 10) for each overlap are shown, ranked by Normalized Enrichment Score. For all graphed pathways, FDR.q < 0.002.

We next asked if these changes in gene expression were represented only in genes strongly up- or down-regulated by MYC. GSEA can detect small differences across many genes [56], so we sought to confirm that these programs arose from highly upregulated or downregulated genes by performing a second independent analysis of differential expression using a different functional enrichment platform, ToppFun [49]. We focused on genes that were at least 1.5-fold up- or down-regulated (**Supplemental Figure 4A**), and found that the three cell lines had 794 genes upregulated 1.5-fold in common, and 824 genes downregulated 1.5 fold in common (**Supplemental Figure 4B,C**). We subjected each of these in-common gene sets to ToppFun pathway enrichment analysis, and obtained highly similar results to GSEA analysis. Upregulated programs included mitochondrial and ribosomal biogenesis (**Supplemental Figure 4D**), while downregulated programs included those related to ECM, collagen, and focal adhesion (**Supplemental Figure 4E**). These results suggested strong commonalities in the MYC-driven transcriptional program between three unrelated cell lines representing two different forms of cancer, and are consistent with and corroborate the role of MYC in up-regulating ribosome and mitochondrial biogenesis as well as down-regulating cell adhesion [29, 63].

### Oncogenic MYC impairs transcriptional oscillation to drive static gene expression programs

Given that MYC disrupts normal molecular circadian clock rhythmicity across multiple systems, a key question arises: which oscillatory programs are shifted by MYC from being oscillatory in MYC-OFF to being statically up- or down-regulated when MYC is activated? The answer to this question would explain how MYC shifts the behavior of cells by disrupting their molecular clock and circadian rhythmicity, and begin to explain what benefit MYC-amplified cancers gain from disruption of the molecular circadian clock. We asked this question both at the pathway-level and at the gene-level using complementary methodologies. At the pathway-level, we first determined which gene expression programs were oscillatory in MYC-OFF cells, as determined by PSEA (as discussed in **Figure 3**), and lost oscillation when MYC was turned on. We then examined overlap between these programs that lost oscillation and which programs were determined to be up- or down-regulated by MYC via GSEA analysis (see schematic in **Figure 5A**). In both SHEP and SKNAS, programs that lost oscillation and became upregulated corresponded to RNA metabolism and ribosomes, as well as cell cycle programs in SKNAS (**Figures 5B-C**). This was similar to those programs that were upregulated by MYC in these cells (see **Figure 4B** and **Supplemental Figure 4B**). Those programs that switched from being oscillatory to downregulated corresponded to endoplasmic reticulum and aging in SHEP, and extracellular matrix, morphogenesis, and plasma membranes in SKNAS (**Figures 4B-C**). These gene sets were overall less similar to those downregulated by MYC in these cell lines (see **Figure 4C** and **Supplemental Figure 4C**). Conversely, we did not detect in U2OS overlap between oscillating programs and those that became up or down-regulated (not shown), suggesting that disruption of oscillation in U2OS may occur more at the protein or metabolite level.

**Figure 5.**
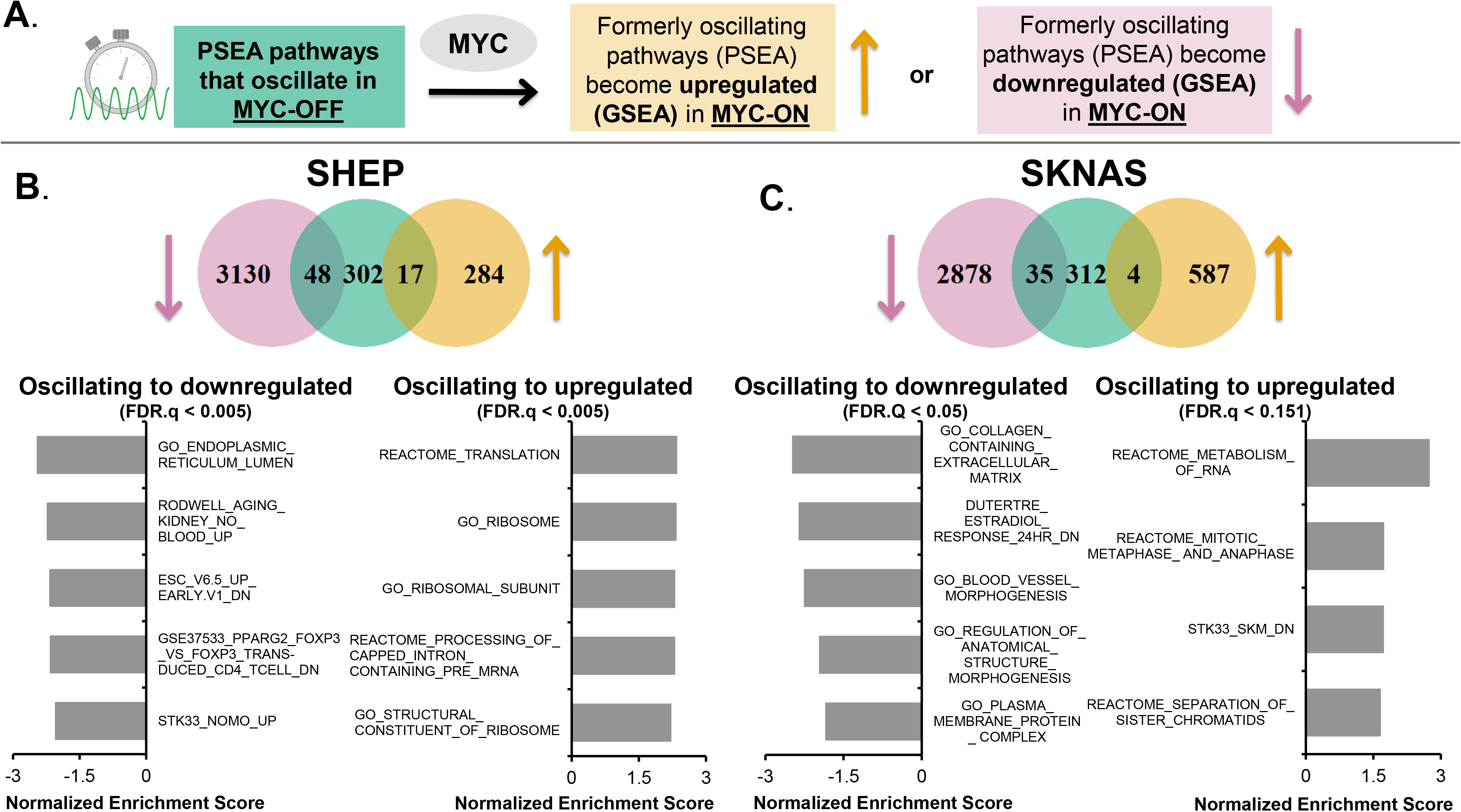
Oncogenic MYC shifts oscillatory gene expression to static non-oscillatory regulation. **A.** Illustration of workflow to identify pathways that lose oscillation when MYC is activated (by PSEA), and which of these pathways become up- or down-regulated by MYC (using GSEA). **B, C**. Venn diagram from SHEP N-N-MYC-ER (**B**) and SKNAS N-MYC-ER (**C**) of pathways (identified by PSEA) that were circadian in MYC-OFF (center, green), and lost oscillation and became either downregulated (left, pink) or upregulated (right, tan) in MYC-ON (identified by GSEA). The most highly significant downregulated and upregulated pathways that lost oscillation from SHEP (**B**) and SKNAS (**C**) are shown below each Venn diagram, as ranked by Normalized Enrichment Score from GSEA. For all data shown, p value is < 0.01 and FDR.Q value is indicated on the graph.

We cross-validated these results with an independent approach that focused on the gene level (**Supplemental Figure 5**). In this approach, we first identified genes that had significant circadian oscillation in MYC-OFF, as identified in ECHO, and lost this oscillation in MYC-ON (see **Figure 2**). Of these genes that lost oscillation in MYC-ON, we next asked which were significantly up- or down-regulated by MYC, using DeSeq2, and finally queried these overlapping genes for gene set enrichment by ToppFun (see schematic in **Supplemental Figure 5A**). In SHEP and SKNAS, programs corresponding to genes that switched from being oscillatory to upregulated were similar to those determined by PSEA and GSEA in **Figure 5**, corresponding to ribosome assembly, tRNA aminoacylation and biosynthesis, mitochondrial activity, cell cycle and protein translation (**Supplemental Figure 5A,B**). For SKNAS, many programs corresponding to the molecular clock also switched from being oscillatory to upregulated (**not shown**). In contrast in U2OS, upregulated programs mostly corresponded to the molecular clock itself as well as macromolecular protein complex programs (**Supplemental Figure 5C**). There were comparatively fewer downregulated programs that were identified with this approach: in SHEP, the few downregulated programs corresponded to cell adhesion / binding and organization, while in U2OS the downregulated programs corresponded to circadian rhythm and cell organization; there were no detected downregulated programs in the queried gene set libraries in SKNAS (**not shown**). Overall, these results suggested that, across multiple cancer models, MYC disrupts circadian transcriptional oscillation and instead promotes static (non-oscillatory) regulation of processes such as biosynthesis, metabolism, and extracellular matrix-related gene expression programs.

### Rhythmic expression of nutrient transporters is disrupted by MYC

Intracellular metabolism is known to be highly circadian [10, 64-66], potentially balancing anabolic and catabolic metabolism in concert with whole body rhythms. In contrast, cancer cells, particularly those driven by MYC, engage in upregulated nutrient uptake and biosynthesis [29, 67]. Our data indicated that in multiple cell lines, MYC caused metabolic and biosynthetic processes that were formerly circadian to flip to static upregulation (**Figure 5, Supplemental Figure 5**). In addition, we have previously observed that MYC expression in U2OS ablated oscillation of intracellular glucose while boosting uptake of glucose and glutamine [31]. Since these processes are regulated in part by nutrient transporter expression and availability, we asked whether nutrient transporter rhythmicity and expression is affected by MYC in three cell models. We focused on the two subunits of the LAT1 amino acid transporter: LAT1 (*SLC7A5)* and 4F2hc (*SLC3A2*, also known as CD98), which transport glutamine and other amino acids, as well as the ubiquitously expressed GLUT1 glucose transporter (*SLC2A1*) [68-70]. GLUT1 and 4F2hc are glycosylated, which aids in their trafficking to the plasma membrane [71, 72], so we performed immunoblot using a technique to determine glycosylation level of these proteins [72]. Where appropriate, two different exposures of the same immunoblot (‘dark’ and ‘light’) are shown. For each cell line, n=3-4 biological replicates were performed, quantified, and analyzed for both changes in total protein level, and for oscillation with ECHO. For all three cell lines, LAT1 oscillated in a circadian fashion in MYC-OFF, and this was lost in MYC-ON (**Figure 6A-C, quantification in bottom panels**). At the same time, LAT1 was significantly upregulated by MYC in all three cell lines (**Supplemental Figure 6**). Glycosylated 4F2hc oscillated in MYC-OFF and lost oscillation in MYC-ON in SKNAS but was upregulated by MYC in all three cell lines (**Figure 6A-C, Supplemental Figure 6**). We also observed that the genes encoding LAT1 and 4F2hc, *SLC7A5* and *SLC3A2*, were significantly upregulated by MYC in all three cell lines (**not shown**). GLUT1 oscillation was not ablated by MYC in any of the cell lines, and it was only upregulated by MYC in SHEP (**Figure 6A-C, Supplemental Figure 6**), in contrast to previous reports that MYC transcriptionally upregulates *SLC2A1* in other cancer models [73]. This suggested a potential specificity for MYC regulation of oscillation and expression of amino acid transport over glucose transport in these cell lines. As has been shown previously [31, 32], we also observed suppression of BMAL1 and upregulation of REV-ERBα regardless of timepoint (**Figures 6A-C, Supplemental Figure 6**). Overall, these data suggested that nutrient transporter expression and glycosylation is rhythmically regulated and ablated by MYC, as MYC increases post-transcriptional modification and levels of these transporters.

**Figure 6.**
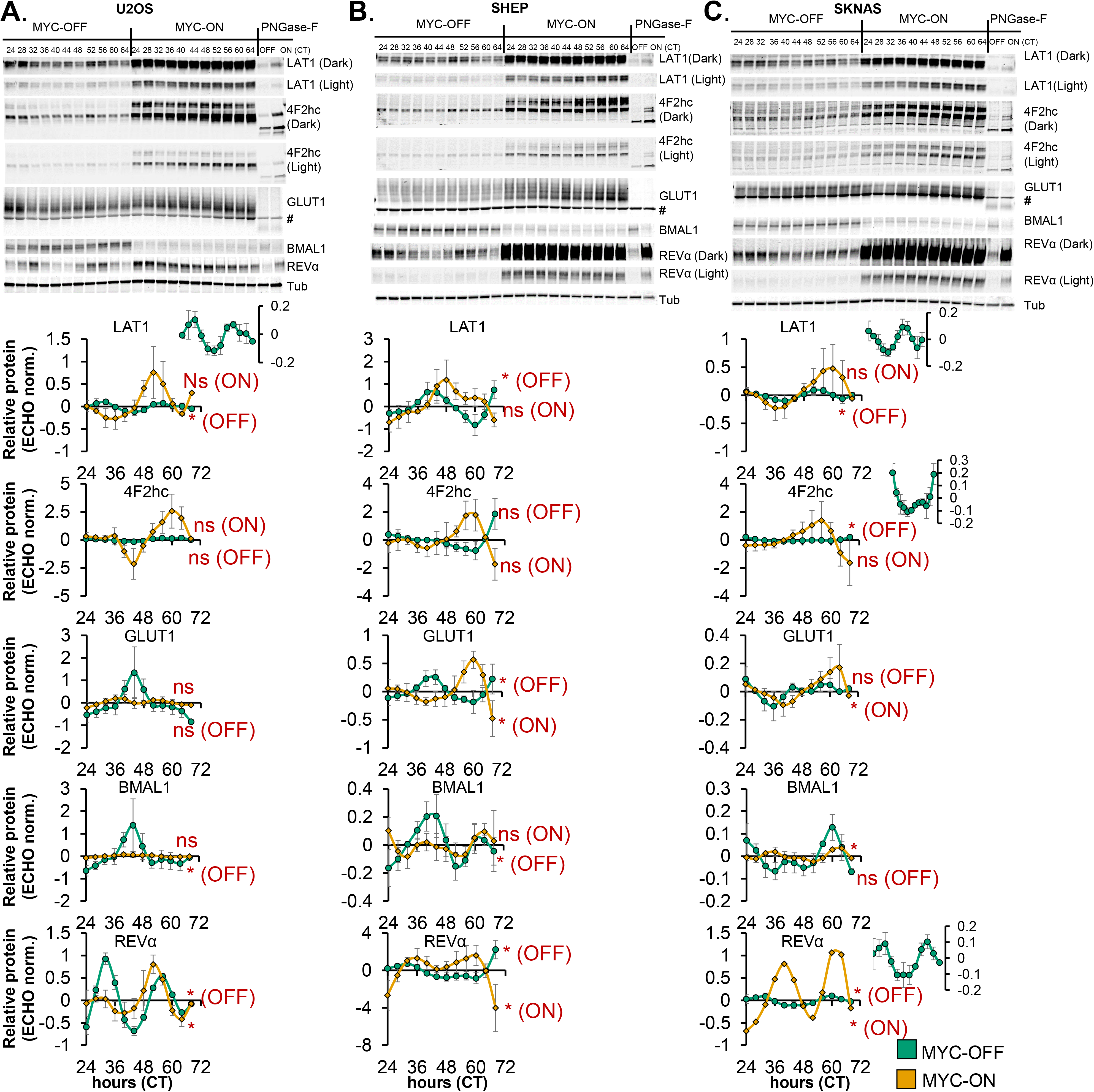
Rhythmic expression of nutrient transporters is disrupted by MYC. A-C, top panels. U2OS MYC-ER (**A**), SHEP N-MYC-ER, (**B**), and SKNAS N-MYC-ER (**C**) were treated with ethanol control (MYC-OFF) or 4-hydroxytamoxifen (MYC-ON) (4OHT) to activate MYC, and entrained with dexamethasone, and after 24 hours, protein was collected every 4 hours for the indicated time period. Protein lysates were prepared to preserve protein glycosylation (see Methods), and immunoblot was performed for the indicated proteins. For some targets [4F2hc, LAT1, REV-ERBα (abbreviated REVα)], a darker exposure (‘dark’) and lighter exposure (‘light’) of the same blot are presented. For GLUT1, # indicates a non-specific band. Some samples (CT26 for U2OS, CT32 for SHEP and SKNAS) were treated with PNGase-F prior to immunoblot to remove glycosylation marks. Note that the PNGase-F lanes have less protein loaded than the other lanes. Data represent n=3-4 biological replicates for each cell line. **A-C, bottom panels.** Results from n=3-4 immunoblot replicates were quantified, relative to Tubulin, and analyzed in ECHO for circadian rhythmicity. Displayed curves are the baseline-subtracted and smoothed outputs from ECHO analysis. ECHO norm. = ECHO normalized. Proteins with a 22-26 hour period and a p value < 0.05 and a BH.Adj.P.Value < 0.05 were deemed rhythmic, which is indicated in the Figure. Note the inset MYC-OFF only graphs for U2OS LAT1, SKNAS LAT1, SKNAS 4F2hc, and SKNAS REV-ERBα, which show oscillation of these proteins in MYC-OFF cells on a different scale.

The above findings on nutrient transporter glycosylation did not demonstrate whether these transporters actually reached the cell surface, whether cell surface expression of transporters was increased by MYC, and whether these resulted in identifiable metabolic changes in the cells. To address this, we performed an On-cell Western [74], which stains fixed and unpermeabilized cells in a dish to determine surface expression of a target protein. We focused on LAT1, and found that surface expression of LAT1 was significantly increased in MYC-ON as compared to MYC-OFF cells (**Figure 7A,B**), suggesting that an increase in expression and glycosylation correlates with more transporter on the cell surface. To further determine whether changes in nutrient transporter expression, glycosylation, and localization correlated with increases in intracellular metabolite pools, we performed ultra-performance liquid chromatography-tandem mass spectrometry (UPLC-MS/MS) metabolomics in U2OS, in the presence or absence of activated MYC. In two independent experiments, we found that almost all amino acids were significantly increased in MYC-ON (**Figure 7C, Supplemental Figure 7**), suggesting that increased nutrient transporter expression and glycosylation correlated with enhanced intracellular amino acid pools. Overall, these data showed that MYC ablates nutrient transporter oscillation and increases transporter expression, cell surface localization, and activity.

**Figure 7.**
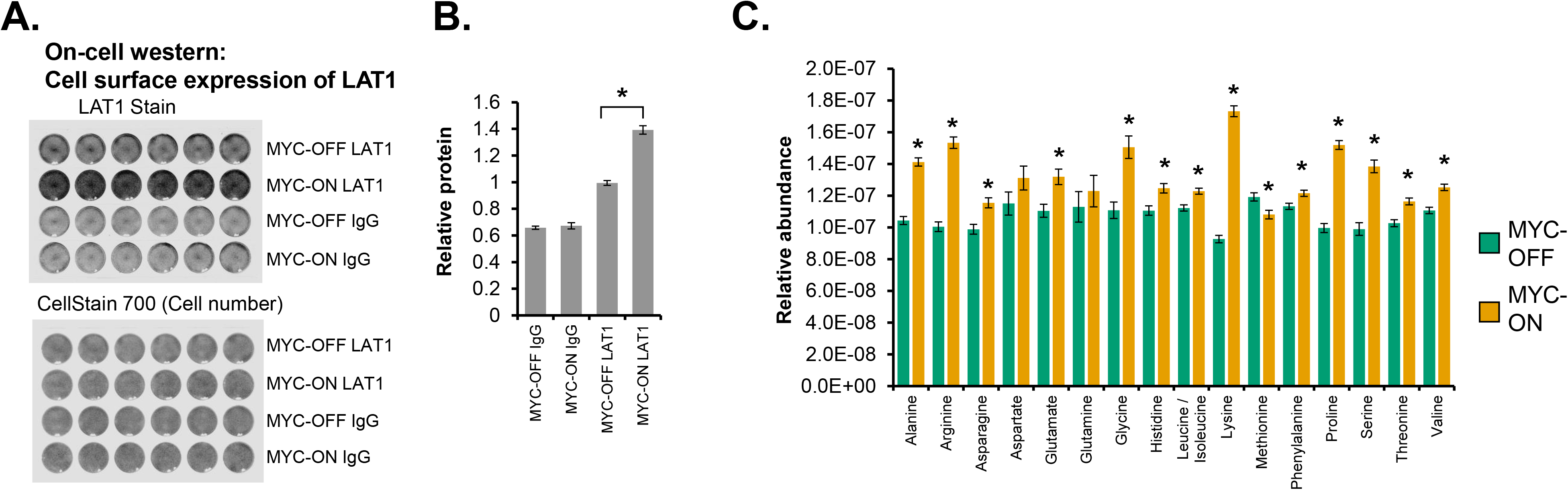
MYC increases LAT1 surface localization and intracellular amino acid pools. **A.** On-cell western of U2OS MYC-ER cells ± MYC for LAT1. U2OS MYC-ER cells were grown on a 24-well tissue culture plate ± 4OHT for 48 hours, fixed with formaldehyde but not permeabilized, and then stained with the indicated antibody or IgG control. CellStain 700 indicates cell density in each well. Data represent at least two independent experiments of 6 biological replicate wells each. **B.** Quantitation of LAT1 or IgG from (**A**). * indicates P < 0.00001 by Welch’s Corrected Student’s T-test. **C.** LC-Mass spectrometry was performed on U2OS MYC-ER treated ± 4OHT for at least 48 hours. N=25 circadian timepoints for MYC-OFF and MYC-ON were averaged as biological replicates, normalized to cell number for each collection. * indicates p < 0.05 by Welch’s Corrected Student’s T-test.

### MYC disrupts metabolic circadian oscillations in a cell-autonomous manner

The ablation of oscillatory nutrient transporter glycosylation and expression, along with our previous analysis of metabolic oscillations by NMR and mass spectrometry [10, 31], raised the intriguing possibility that MYC may disrupt global cell-autonomous metabolic oscillation. To test this, we performed time-series metabolomic analysis using UPLC-MS/MS at 2-hour intervals in dexamethasone-entrained U2OS MYC-ER cells over two independent experiments, termed Replicate A and Replicate B. Using the ECHO algorithm, we found in Replicate A that there were 37 oscillating metabolites in MYC-OFF, while only 6 of these metabolites remained oscillatory with a ∼24-hour period in MYC-ON (**Figure 8A**). The traces of individual metabolites were plotted for MYC-OFF and MYC-ON in **Figure 8B**. Similar results were observed in Replicate B (**Supplemental Figure 8A,B**). In concordance with our transcriptomic observations, some metabolites in both Replicates gained 24-hour rhythmicity in MYC-ON that did not previously have this rhythmicity in MYC-OFF (**Supplemental Figure 8C-D**).

**Figure 8.**
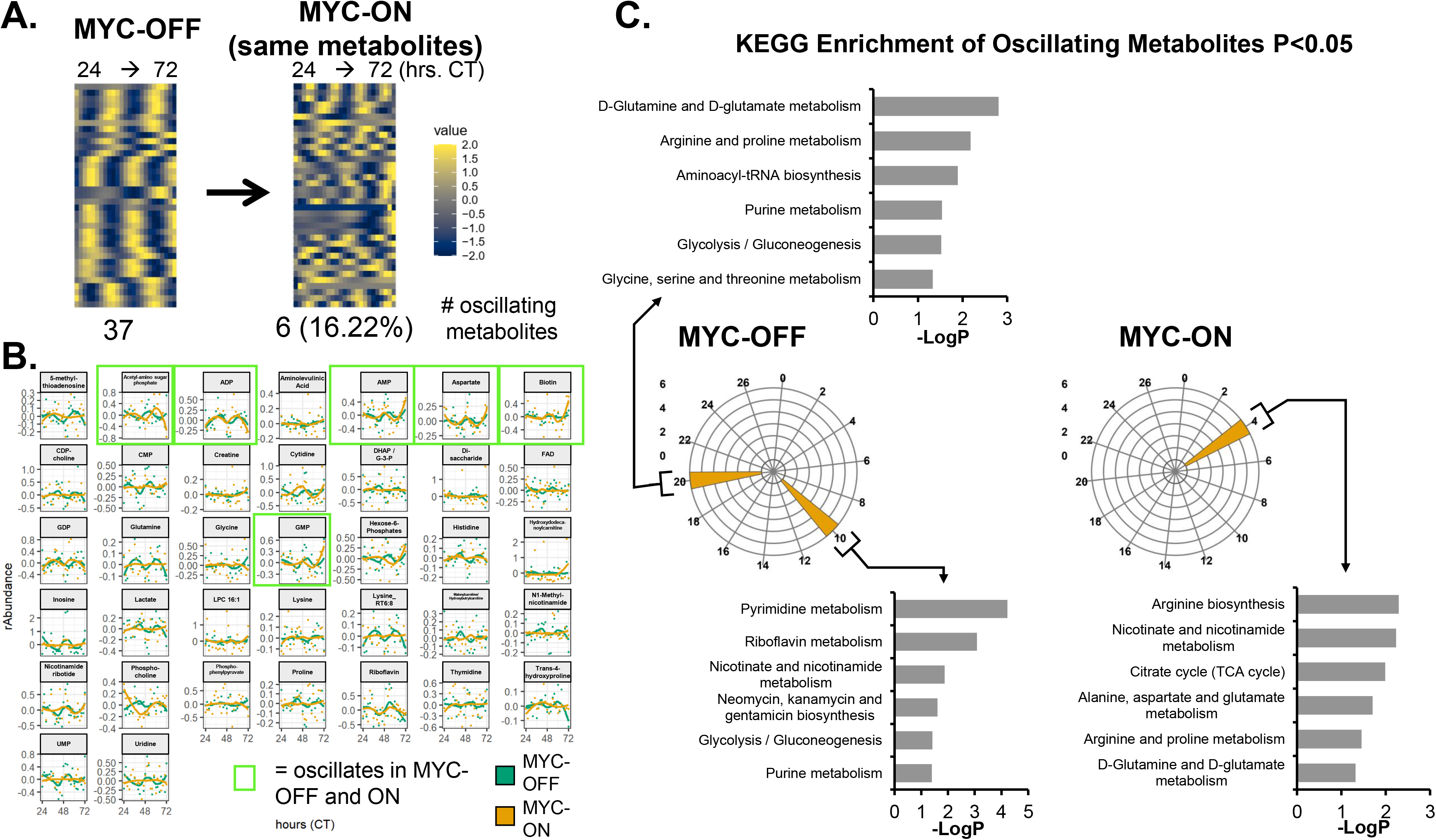
MYC disrupts metabolic circadian oscillations in a cell-autonomous manner. U2OS MYC-ER were treated with ethanol control (MYC-OFF) or 4-hydroxytamoxifen (MYC-ON) (4OHT) to activate MYC for 24 hours, and entrained with dexamethasone, and after 24 hours, cells were extracted for polar metabolites every 2 hours for the indicated time period. Mass spectrometry was performed, and rhythmicity was assessed by ECHO for both MYC-OFF and MYC-ON, with metabolites with a 20-28 hour period and with BH.Adj.P.Value < 0.05 deemed rhythmic. These metabolites were sorted by phase and are presented in a heatmap for MYC-OFF. For MYC-ON, the same metabolites that are rhythmic in MYC-OFF are presented in the same order, but with MYC-ON values instead. **B.** Metabolites deemed to be oscillating by ECHO for MYC-OFF are graphed, with dots representing the relative abundance values, and lines indicating the fitted oscillation curves as calculated by ECHO. Metabolites with a green border are oscillatory in both MYC-OFF and MYC-ON. **C.** KEGG enrichment analysis was performed on metabolites that peaked in the indicated phases in MYC-OFF or MYC-ON conditions, and significantly enriched pathways were graphed on a polar histogram. For each histogram, the scale is on the left side. Pathways with a p < 0.05 were deemed significant and are graphed.

Finally, we queried which metabolic programs were circadian in MYC-OFF and MYC-ON by performing KEGG enrichment analysis. Peak phase of metabolites was determined, KEGG analysis was performed separately on each set of oscillating metabolites, and the resulting significantly enriched pathways were plotted in polar histograms. In both Replicates A and B for MYC-OFF, we observed peaks of oscillation at CT10 and CT20. Those at CT10 corresponded to programs in nucleotide metabolism, while those at ZT20 instead were enriched for programs in amino acid metabolism and tRNA biosynthesis (**Figure 8C and Supplemental Figure 9A**). This suggested that in cells without overexpressed MYC, metabolite programs oscillate in a predictable and consistent pattern. In contrast, there was no discernable pattern in MYC-ON cells: in Replicate A, metabolic programs were clustered around a single timepoint, while in Replicate B, they were fragmented in a manner distinct from MYC-OFF (**Figure 8C, Supplemental Figure 9B**). These observations demonstrate that cells without amplified ectopic MYC engage in circadian oscillatory metabolism that temporally separates different bioenergetic processes, which is disrupted when MYC is elevated and active.

## Discussion

We and others previously showed that MYC disrupts molecular clock oscillation in cancer cells, but the reasons for this, and what potential benefit cancer cells might gain, remained unclear. Our findings, in sum, indicate that overexpressed MYC suppresses circadian oscillation of transcriptional and metabolic programs across multiple models of cancer (**Figure 9**). MYC overexpression led to a greater than 85% loss of global transcriptional oscillation in neuroblastoma and osteosarcoma cell lines (**Figure 2A**). To identify robust signatures of which genes were circadian in MYC-OFF (low-MYC) cell lines, and which genes were regulated by MYC in MYC-ON (overexpressed MYC) cell lines, we focused on genes and pathways that overlapped between at least two of the three cell lines in our study. In the absence of overexpressed MYC, we found that oscillating genes which overlapped between multiple cell lines were enriched for pathways associated with cell cycle / DNA replication, endoplasmic reticulum, and cell junctions / adhesion (**Figure 3**). When MYC was activated, we found that MYC upregulated genes associated with metabolism and biosynthesis, and suppressed genes associated with adhesion and extracellular matrix (ECM) (**Figure 4 and Supplemental Figure 4**). Our observation that MYC globally downregulates genes and programs associated with adhesion and ECM substantiates and adds to the finding of MYC in suppressing keratinocyte cell adhesion [63]. More importantly, we identified, that genes and pathways that oscillate in the absence of overexpressed MYC lose oscillation when MYC is activated and instead become up- or down-regulated. Notably, metabolic and biosynthetic processes were most significantly enriched in pathways and genes that shifted from oscillatory in MYC-OFF to upregulated in MYC-ON (**Figure 5 and Supplemental Figure 5**). These results indicated that cells with oncogenic MYC may release metabolic genes from circadian control, instead upregulating metabolic gene expression programs without regards to oscillation. Further focusing on metabolism, we found that the LAT1 amino acid transporter (constituting LAT1 and the 4F2hc subunit) demonstrated oscillatory expression in MYC-OFF, and this rhythmic oscillation was lost in MYC-ON, while MYC massively upregulated LAT1 expression across multiple cell lines (**Figure 6 and Supplemental Figure 6**). We also found that LAT1 surface localization was enhanced by MYC (**Figure 7A**). Interestingly, this was specific to the LAT1 transporter, as MYC did not consistently alter GLUT1 oscillation or expression across cell lines (**Figure 6**). Finally, oscillation of metabolites themselves were also suppressed by MYC, and circadian segregation of anabolic and catabolic metabolic processes was ablated (**Figure 8 and Supplemental Figure 9**). Overall, our findings for the first time show MYC suppresses oscillation of genes associated with metabolism and biosynthesis to instead upregulate these processes and engage in heightened metabolism in an oscillation-independent fashion.

**Figure 9.**
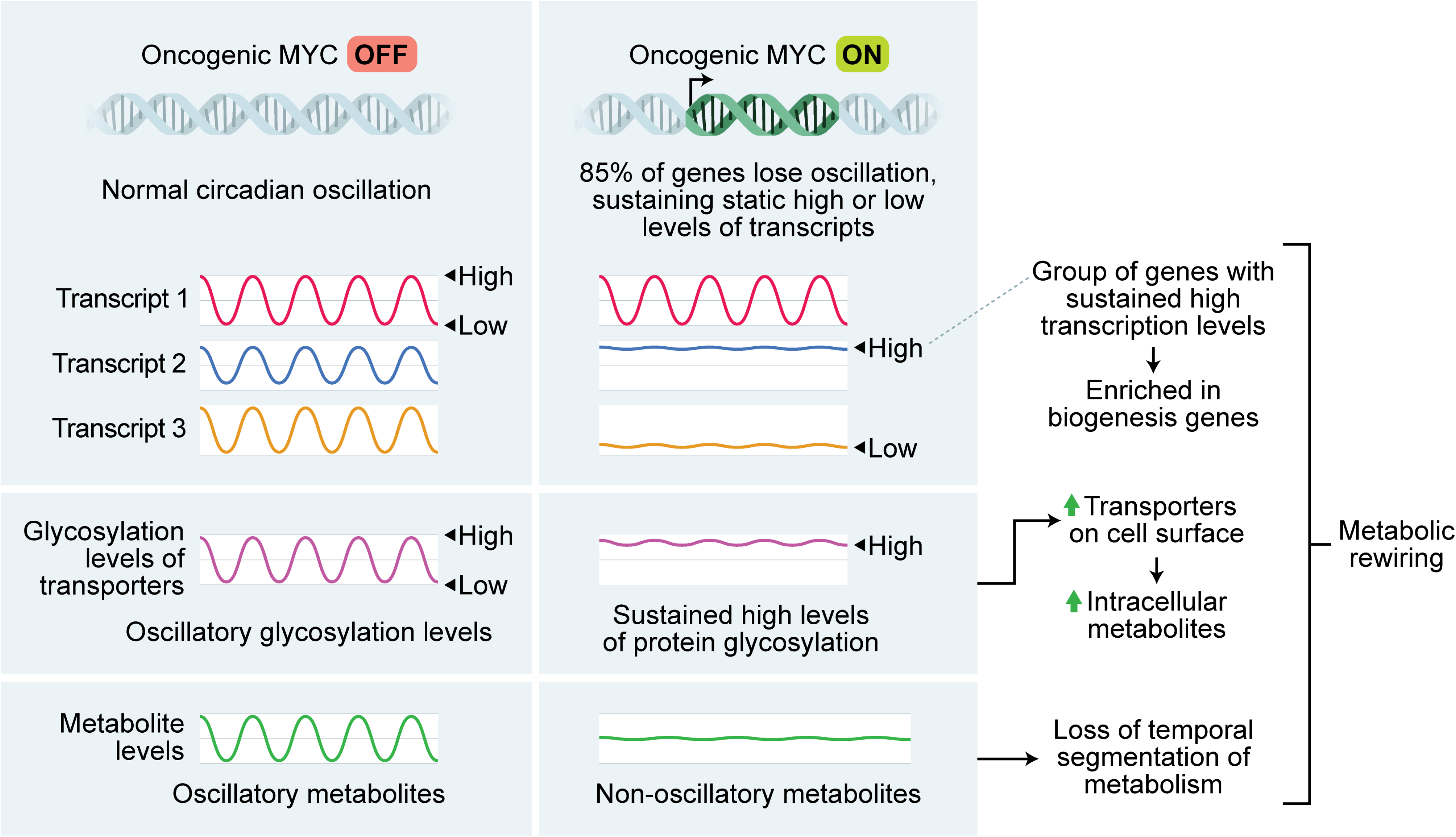
Graphical Abstract.

Our findings that MYC shifts certain metabolic and cell-attachment pathways from being oscillatory to being up- or down-regulated is supported by observations of similar phenomena from human cancer samples. It has historically been difficult to assess the state of the molecular circadian clock in human tumor samples since these samples are from single timepoints, and time of day information is often unavailable. However, several recent studies using distinct methodologies have all determined that rhythmicity tends to be dampened or ablated in human tumors compared to normal tissue [22-24, 75]. One weakness of these techniques is that, currently, they cannot detect phase or period changes in oscillation, only amplitude changes. Nonetheless, in addition to identifying lost rhythmicity of the molecular clock in tumors, these studies interrogated which pathways had decreased oscillation in tumors. One study, using a methodology known as LTM (Lunch Table Method), found across 11 tumor types that the pathways most associated with circadian rhythmicity in tumors were those related to extracellular matrix, collagen, and TCA cycle and electron transport (metabolism) [75]. In our study, we observed that these same pathways in MYC-ON cells were among the ones most likely to lose oscillation and become upregulated (metabolism) or suppressed (ECM and collagen) (**Figure 5 and Supplemental Figure 5**). The authors of the LTM study speculated that the correlation between ECM / collagen rhythmicity and tumor rhythmicity may be related to cellularity of the tumor (ie, tumors with more endothelial and fibroblast cells have less cancer cells and more rhythmicity) [75], whereas our findings suggest a possible tumor-cell autonomous mechanism for loss of ECM rhythmicity in cancer cells. A separate study, using a methodology known as CYCLOPS (cyclic ordering by periodic structure), found that pathways related to redox metabolism and hypoxia were dampened in hepatocellular carcinoma (HCC) compared to normal liver [22]. More intriguingly, the authors also identified GLUT2 (*SLC2A2*) as a transcript that displayed dampened oscillation in HCC compared to normal liver and kidney. These findings are similar to our observations that MYC upregulates and ablates oscillation of nutrient transporters and metabolic pathways across multiple cancer types. The authors of the CYCLOPS study did not identify a mechanism for this loss of oscillation, but since HCC is known to be driven by amplified MYC [76], it is possible to speculate that elevated and deregulated MYC in HCC may drive this loss of rhythmicity in metabolic processes. Interestingly, in U2OS where our metabolomics studies were conducted, we observe loss of oscillation of the LAT1 nutrient transporter when MYC is activated (**Figure 6**), but this does not directly tie back to loss of oscillation of metabolic gene expression programs (**Figure 5, Supplemental Figure 5**). This may be due to the fact that some protein and metabolic circadian oscillations arise in a manner independent of direct transcriptional control [8-10]. Nonetheless, our findings for the provide a potential mechanism for the loss of oscillating pathways and metabolic processes in cancer cells.

MYC is often overexpressed in cancer, and in many cases, this overexpression arises from either genomic translocation or amplification, which occur in at least 9-28% of all human cancers [27]. The MYC-ER and N-MYC-ER systems model genomic amplification in cancer, where MYC is no longer under the control of endogenous promoter and other regulatory elements [77]. In these cancer models where MYC is amplified, which represent osteosarcoma and neurobalstoma, we and others have shown that MYC suppresses or interferes with BMAL1 and disrupts molecular clock oscillation [31-36]. Our results now show that when MYC disrupts the molecular clock, this is accompanied by widespread suppression of genetic and metabolic oscillations that occur in the absence of amplified MYC (**Figures 2 and 8**). More notably, our findings suggest that MYC “releases” biosynthetic and metabolic processes from circadian control in order to be upregulated without regards to oscillation (**Figure 5 and Supplemental Figure 5**).

The molecular circadian clock is not disrupted in all cancers, and in particular seems to be maintained (albeit in an altered state) in those driven by mutated HRAS or KRAS [78-80]. In fact, alteration (but not complete ablation) of the circadian clock may enhance MYC expression even in cases where MYC is not amplified, and MYC itself is circadian-regulated and oscillatory in these settings [12, 17, 43, 81, 82]. In cancer models where MYC is upregulated (by another oncogene or through circadian disruption), and is still circadian-controlled, MYC does not seem to exert a strong effect in suppressing BMAL1 and ablating rhythmicity of the molecular clock or downstream targets [11, 17, 43, 81, 82]. One notable exception is in glioblastoma stem cells (GSCs), where GSCs with amplified MYC maintained oscillation and actually relied on CLOCK and BMAL1 to maintain stemness [83]. A similar necessity for the circadian clock was seen in acute myeloid leukemia stem cells [84]. It may be that GSCs (and other select cancer stem cell populations) with amplified MYC specifically select for rare populations where clock function is somehow maintained. Additional studies are necessary to further define the context of MYC and maintenance of the core circadian clock in tumors that are believed to be driven largely by the cancer stem cell compartment.

We observed that some genes and pathways gained oscillation when MYC was activated (**Supplemental Figures 2 and 3**). Generally, fewer pathways were enriched in MYC-ON oscillatory genes than MYC-OFF oscillatory genes, especially when using the ToppFun time-independent approach, indicating some degree of stochasticity in which genes gained oscillation. This was particularly true in SKNAS, where only two gene sets were enriched in MYC-ON in our PSEA analysis (**Supplemental Figure 3A**). Perhaps not coincidentally, SKNAS also had the highest degree of *MYC* or *MYCN* overexpression of the three cell lines used in this work (**Supplemental Figure 1A**). One notable group of genes that gained oscillation were cell cycle-related genes in SHEP and U2OS (**Table 1 and Supplemental Figure 3**), consistent with the notion that MYC induces cell cycle advance [28]. Interestingly, several recent papers reported that metabolic oscillations associate with active cell cycle, though there was not consensus on whether these oscillations depended on cell cycle progression [85, 86]. Thus, it is possible that the metabolic oscillations we observe in MYC-ON (**Figure 8, Supplemental Figures 8-9**) originate from cell cycle progression and not from the molecular clock. Nonetheless, the source of gene and metabolite oscillations in MYC-ON cells remains unclear. We note that while BMAL1 is suppressed by MYC, its expression is not ablated (**Figures 1,6** and [31-34, 36]). Thus, it is possible that these oscillations originate from residual CLOCK-BMAL1 activity. Indeed, we previously showed that MYC competes with BMAL1 for binding to the *NR1D1* (REV-ERBα) promoter [31]. This is similar to a previously published mechanism whereby the transcription factor USF1 competes with CLOCK-BMAL1 for DNA binding and may alter outputs of the molecular clock [87]. We speculate that MYC, when amplified, may similarly compete for CLOCK-BMAL1 binding sites genome-wide, and oscillations that emerge in MYC-ON cells may arise from new sites occupied by CLOCK-BMAL1 in circumstances where MYC competes for their normal binding sites. It is notable that major perturbations to the molecular clock machinery, such as deletion of REV-ERBα and β, can cause a major remodeling of oscillation transcriptional output and the emergence of many new oscillating genes, as was recently shown in the liver [88]. A future line of inquiry should determine if non-canonical circadian oscillations that emerge in MYC-amplified cancer cells contribute to proliferation or metabolic phenotype.

Metabolic rewiring supports cancer cell proliferation and growth, and our findings suggest that loss of circadian oscillation by MYC may be a feature of metabolic rewiring. This rewiring relies not only on the aberrant expression and activity of enzymes from the glycolytic, oxidative, and biosynthetic pathways but also on optimizing the influx and efflux of nutrients by solute carrier proteins (SLCs) [67]. Notably, all these processes are regulated by oncogenic MYC [29, 89]. One example is the loss of rhythmicity we observed in the LAT1 amino acid transporter. LAT1 a heterodimeric transporter complex of LAT1 (*SLC7A5*) with 4F2hc (*SLC3A2*), which is required for LAT1 functional activity, stability, and proper location in the plasma membrane [68]. LAT1 is a key mediator of essential amino acid uptake, promoting amino acid homeostasis and mTORC1 pathway activation in cancer cells, which supports cell proliferation and survival [90, 91]. The LAT1 subunit is upregulated in various cancers, while its inhibition reduces tumor growth [91, 92]. Our results show that MYC activation ablates oscillation of the LAT1 subunit and increases LAT1 total protein expression and membrane localization (**Figure 6, Supplemental Figure 6 and Figure 7**), corroborating previous findings of MYC-dependent upregulation of LAT1 in cancer and T cells [92-96]. MYC effects on LAT1 may rely on transcriptional mechanisms, since MYC binds to E-box regions in the LAT1 promoter [93]. LAT1 forms a heterodimeric transporter complex with 4F2hc, also a MYC target, which is required for LAT1 functional activity, stability, and proper location in the plasma membrane [68, 97, 98]. 4F2hc harbors four N-glycosylation sites that are required for proper stability and trafficking to the membrane. Mutation of those sites correlates with a lower abundance of LAT1 in the plasma membrane and reduced transport activity [71]. We found in our cancer cell modes that when MYC was activated, increased expression of glycosylated 4F2hc was detected in all three cell lines, and oscillation of 4F2hc was lost in SKNAS (**Figure 6**). The surface expression of LAT1 and intracellular amino acid pools were significantly increased in MYC-ON compared to MYC-OFF cells (**Figure 7**) in U2OS cells, suggesting that MYC disrupts expression and glycosylation oscillations to drive static but highly active transporter states.

Overall, our data bring new mechanistic insights into how MYC disrupts cell-autonomous metabolic circadian oscillation. We speculate that in the MYC-OFF state, the circadian oscillatory expression of transporters and metabolic enzymes as well as the sub-cellular trafficking of metabolite transporters allows for cell metabolism to synchronize with organismal circadian feeding and fasting cycles. In the MYC-ON state, mimicking MYC amplification, we surmise that induced expression of transporters and enzymes is relieved from circadian for heightened anabolic metabolism to support oncogenic cell growth. It remains to be fully determined, however, how this loss of circadian control of metabolism and gain of heightened metabolic flux in MYC-amplified tumor cells may directly contribute to tumorigenesis. In this respect, it is notable that increased glutamine flux with MYC activation in the human lymphoma and mouse liver cancer models is associated with increased tumorigenesis that can be attenuated by blocking glutamine utilization either genetically or pharmacologically [99-101]. Additionally, increased metabolic fluxes by MYC may support oscillations tied to cell cycle or other rhythms to become more predominant and better drive tumor cell growth [85, 86, 102, 103].

Our findings have implications for potential applications of chronotherapy in the treatment of cancer. Chronotherapy is defined as timed administration of a drug or treatment based on circadian information, to maximize efficacy and / or minimize toxicity [104]. Given that the majority of drug targets are likely to be circadian [4], chronotherapy has much promise in cancer biology in particular. However, the question of approach remains: should a chronotherapy strategy be designed to target the tumor at a particular time when it would be most vulnerable, or instead designed to minimize toxicity? These two approaches may be mutually exclusive. Our findings suggest that circadian phenotype of the tumor matters in this question. For those tumors with MYC amplification, drug targets related to metabolism and biosynthesis may lose rhythmicity, which would favor a chronotherapy approach designed to reduce toxicity. Indeed, the Authors of the CYCLOPS study noted that since GLUT2 is only weakly rhythmic in HCC, a cancer often driven by MYC, the use of the GLUT2-targeting streptozocin should be timed to minimize dose-limiting toxicity in the liver and kidney [22]. In contrast, a separate study focusing on molecular clock gene expression in cancer found that many clinically actionable genes, including those related to metabolism, correlated with clock gene expression across cancer types [21]. This suggests that in cancers that retain rhythmicity (such as those driven by RAS mutations), a chronotherapy approach might instead focus on timing treatment for maximum efficacy against targets in the cancer cells themselves. Overall, our findings suggest that MYC amplification, with further study, might emerge as a prognostic indicator for circadian function which could guide chronotherapy treatment options in the future.

### Limitations of Study

Several limitations of this study are noted. Cancer cells are continuously proliferating, which makes deconvoluting circadian rhythms and cell cycle oscillations challenging. While we did note that some cell cycle genes and pathways shift to oscillatory expression when MYC was expressed (**Table 1, Supplemental Figure 3**), a system where MYC induces proliferation of quiescent cells would be better suited to determine how MYC induces cell cycle rhythmicity. All studies were performed *in vitro* in cell culture conditions. While these conditions are ideal to determine free-running and cell-autonomous oscillations, particularly of metabolites [10, 64, 66], it remains to be determined which oscillatory genes and metabolites MYC disrupts *in vivo*. Finally, we noted that many genes and metabolites gained oscillation when MYC was activated, but did not probe the mechanism of these oscillations (**Supplemental Figures 2,3,8,9**). Since BMAL1 is suppressed but not eliminated by MYC, it is conceivable that these oscillations arise from residual activity of the molecular clock, blocked from its normal activity by elevated MYC levels. Alternately, these oscillations may arise from active cell cycling or in response to gain in oscillation of specific metabolites with MYC-ON (**Supplemental Figures 8,9**).

## Methods and materials

### Cell culture and circadian entrainment

U2OS MYC-ER, SHEP N-MYC-ER, and SKNAS N-MYC-ER were described previously [31, 105, 106]. Cells were cultured in DMEM high glucose (Gibco and Corning) with penicillin-streptomycin (Gibco) and 10% FBS (Hyclone). U2OS MYC-ER cells were cultured with 100 µg/mL Zeocin (Thermo Scientific) except during experiments, to maintain MYC-ER expression. For circadian time-series experiments, cells were plated at 100,000 cells / mL. 24 hours later, cells were treated with ethanol or 0.5 µM 4-hydroxytamoxifen (4OHT) to activate MYC-ER or N-MYC-ER. 24 hours after ethanol or 4OHT treatment, cells were treated with 0.1 µM dexamethasone (Sigma) to entrain the molecular circadian clock. 24 hours after dexamethasone treatment, cells were detached from the plate with trypsin EDTA 0.25% (Gibco) and were collected for RNA in the following intervals: U2OS replicate 1: every 4 hours for 48 hours; U2OS replicate 2: every 2 hours for 48 hours; SHEP replicates 1 and 2: every 4 hours for 52 hours; SKNAS replicate 1: every 4 hours for 52 hours for MYC-OFF, 48 hours for MYC-ON; SKNAS replicate 2: every 4 hours for 48 hours.

### RNA collection and qPCR

Cells were lysed and RNA was isolated using the RNEasy Plus mini kit (Qiagen) or the E.Z.N.A.® HP Total RNA Kit (Omega BioTek). RNA was reverse transcribed to cDNA using the ABI Reverse Transcription Reagents system, using oligo dT for priming (Thermo Scientific). qPCR was performed with cDNA using Power Sybr Green Master Mix (Thermo Scientific) and with the Viia7 quantitative PCR machine (Applied Biosystems). Triplicate technical replicates were performed, outlier replicates (defined as being more than 1 Ct away from other two replicates) were discarded, and relative mRNA was assessed by the ΔΔCt, scaled to the first MYC-OFF timepoint and normalized to *β2M* (β2 microglobulin). Error bars for qPCR experiments are standard error of the mean (S.E.M.). For U2OS qPCR, only the 4-hour resolution timepoints were used for Replicate 2.

### Table of primer sequences used

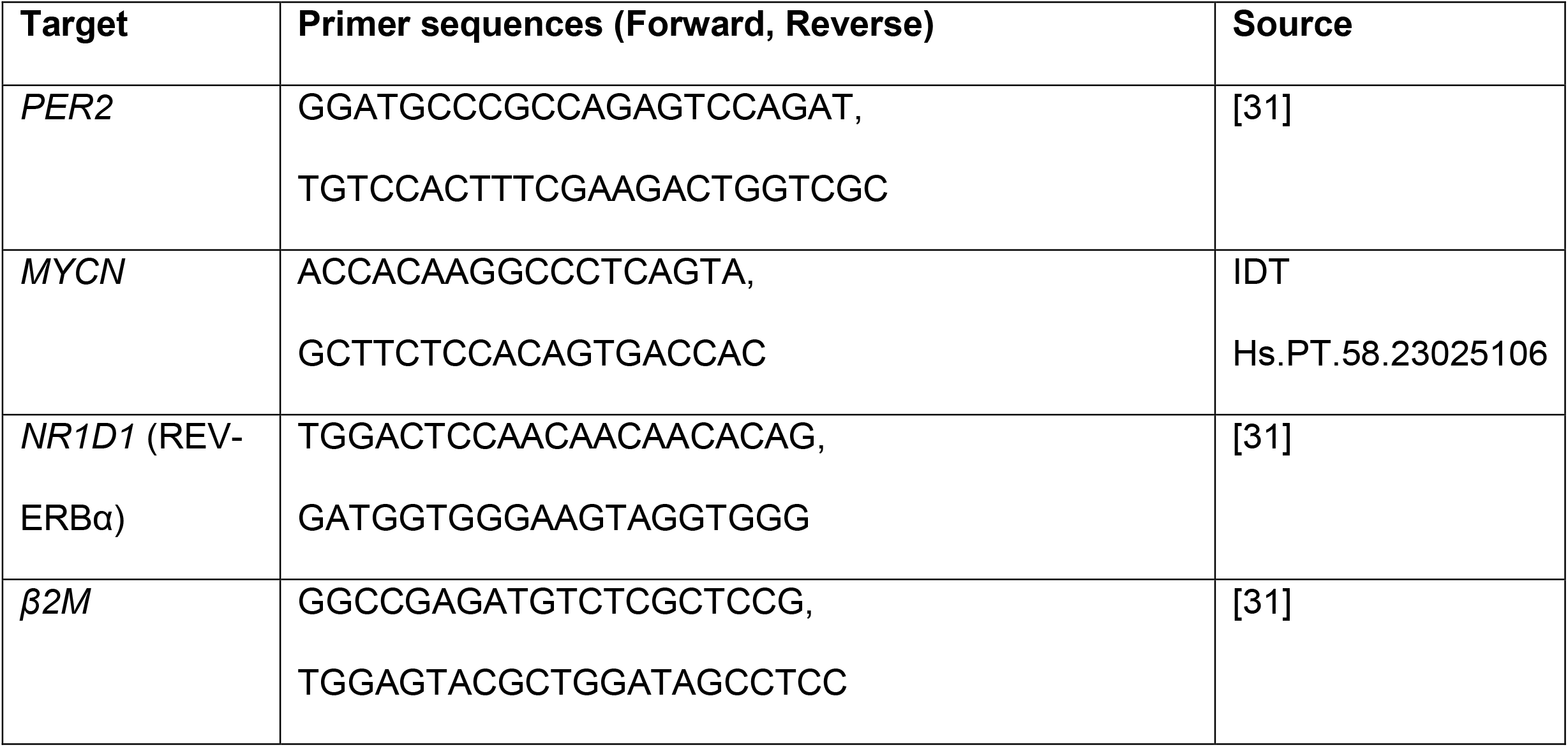

### Protein collection, immunoblot, glycosylation detection, and image quantitation

U2OS MYC-ER, SHEP N-MYC-ER, and SKNAS N-MYC-ER were plated at 100,000 cells / mL. 24 hours later, cells were treated with ethanol or 4-hydroxytamoxifen to activate MYC-ER or N-MYC-ER. 24 hours later, cells were treated with 0.1 µM dexamethasone (Sigma) to entrain the molecular circadian clock. 24 hours after dexamethasone treatment, cells were detached from the plate with trypsin EDTA 0.25% (Gibco) and were collected every 2-4 hours for up to 48 hours. Protein was lysed using the M-Per lysis reagent (Thermo Scientific) with protease inhibitor cocktail (Promega) and phosphatase inhibitors 2 and 3 (Sigma). Lysates incubated on ice for at least 20 minutes, then centrifuged at > 13,000 xg, and supernatant was collected. Protein was quantified using the Bio-Rad DC Protein Assay Kit (Bio-Rad), and lysates of equal concentration were prepared for immunoblot. To detect glycosylated protein, protein samples were prepared with sample buffer that contains 10% glycerol (Sigma) and 5% 2-mercaptoethanol (Sigma), and incubated at room temperature instead of boiled, as previously described [72]. 20 µg of each sample was prepared and run by SDS-PAGE on Bio-Rad Criterion 4-15% 26-well gradient gel (Bio-Rad). For some samples (U2OS CT26, SHEP CT32, SKNAS CT 32), 20 µg was treated with PNGase-F to remove glycosylation marks, according to manufacturer’s instructions (New England Biolabs). For these samples, only 12 µg of protein lysate was loaded onto the Criterion gel. Gels were transferred using the iBlot2 semi-dry blotting system (Thermo Scientific) or Trans-Blot Turbo system (Bio-Rad) to nitrocellulose membranes (Thermo Scientific). The following primary antibodies were used: rabbit anti-4F2hc/CD98 D6O3P mAb (Cell Signaling 13180), rabbit anti-LAT1 (Cell Signaling 5347), rabbit anti-GLUT1 EPR3915 mAb (Abcam ab115730), rabbit anti-REV-ERBα E1Y6D mAb (Cell Signaling 13418), rabbit anti-BMAL1 D2L7G mAb (Cell Signaling 14020) and mouse anti-α-Tubulin DM1A mAb (EMD Millipore CP06-100UG). The following secondary antibodies were used: goat anti-rabbit Alexa Flour 680 (Thermo Scientific A21109) and goat anti-mouse Alexa Flour 790 (Thermo Scientific A11357). Membranes were digitally imaged using a Licor Odyssey CLx infrared imager (Licor) or ChemiDoc MP (Bio-Rad) and uniformly contrasted. Images obtained using the Licor imaging system were quantified with the Image Studio Lite v5.2 Software, which calculated the signal for each band by automatically subtracting the top/bottom average background from the total band signal. Images obtained using the Biorad imaging system were quantified with the Image Lab 6.0.1 software, which calculated the signal for each band by automatically subtracting the background from the total band signal. α-Tubulin was used as the loading control and for normalization.

### On-cell western

On-cell western [74] was performed with U2OS MYC-ER cells. 72,000 cells were plated in each well of a 24-well plate, and 24 hours later, treated ± 4OHT for 48 hours. Cells were then fixed with 3.7% paraformaldehyde (Sigma) but not permeabilized. Wells were washed with tris-buffered saline, and stained with primary antibodies mouse anti-LAT1 BU53 (Novus NBP2-50465AF647) or mouse IgG2A isotype control (Novus IC003R), and secondary antibodies goat anti-mouse Alexa Flour 790 (Thermo Scientific A11357) or CellTag 700 Stain (Licor 926-41090), which is used to quantify total cell number and intensity. The stained plate was then digitally imaged using a Licor Odyssey CLx infrared imager (Licor), and well intensities in both channels (700 nM for CellTag, 800 for LAT1 or isotype control) were quantified using ImageStudio software (Licor). Individual wells were treated as biological replicates, and results are representative of two separate experiments.

### RNA sequencing

RNA-sequencing was performed either at the University of Pennsylvania Genomic and Sequencing Core (U2OS, both replicates), Novogene (SHEP, both replicates, and SKNAS, replicate 1) or the University of Rochester Genomics Research Center (SKNAS, replicate 2). For U2OS, we activated MYC-ER in cells for 24 hours with 4-hydroxytamoxifen (4OHT, MYC-ON), or used vehicle control (MYC-OFF). We then entrained cells circadian rhythms with dexamethasone. 24 hours after dexamethasone treatment, cells were then collected every 4 hours (replicate 1) or 2 hours (replicate 2) for up to 48 hours, and RNA was extracted. CT refers to circadian time, ie, hours after dexamethasone entrainment. RNA at approximately 100 ng/µL was submitted to the University of Pennsylvania Next Generation Sequencing Core, was analyzed by BioAnalyzer and determined to have an average RNA Integrity Number (RIN) of 9.9. RNA-sequencing was performed on ribosome-depleted total RNA (replicate 1) or polyadenylated mRNAs (replicate 2) as 100 base single-end sequencing using an Illumina HiSeq 4000, yielding an average of 74.3 million reads per sample for replicate 1, and 40.9 million reads per sample for replicate 2. For SHEP (both replicates) and SKNAS (replicate 1), we activated N-MYC-ER in cells for 24 hours with 4-hydroxytamoxifen (4OHT, MYC-ON), or used vehicle control (MYC-OFF). We then entrained cells circadian rhythms with dexamethasone. 24 hours after dexamethasone treatment, cells were then collected every 4 hours for up to 52 hours, and RNA was extracted. RNA at approximately 37 ng/µL was submitted to Novogene Co., was analyzed by BioAnalyzer and determined to have an average RNA Integrity Number (RIN) of 9.8. RNA-sequencing was performed on polyadenylated mRNAs using an Illumina NovaSeq 6000 as 150 base paired end sequencing, yielding an average of 54.7 million reads per sample (when adding together both paired ends). For SKNAS replicate 2, we activated N-MYC-ER in cells for 24 hours with 4-hydroxytamoxifen (4OHT, MYC-ON), or used vehicle control (MYC-OFF). We then entrained cells circadian rhythms with dexamethasone. 24 hours after dexamethasone treatment, cells were then collected every 4 hours for up to 48 hours, and RNA was extracted. RNA at approximately 66.67 ng/µL was submitted to the University of Rochester Genomics Research Center (GRC) and was analyzed by TapeStation and determined to have an average RNA Integrity Number (RIN) of 9.3. RNA-sequencing was performed on polyadenylated mRNAs using an Illumina NovaSeq 6000 as 100 base paired end sequencing, yielding an average of 93.1 million reads per sample (when adding together both paired ends). Two biological replicate time-series experiments were performed in each of the three cell lines (SHEP, SKNAS, U2OS).

### Downloading of publicly available RNA-sequencing data: MEF circadian timecourse, macrophage circadian timecourse, and PC3 cells treated with MYCi361

FASTQ files corresponding to a previously published circadian time-series of entrained MEFs in biological triplicate were downloaded from NCBI GEO GSE89018 [43]. FASTQ files corresponding to a previously published experiment of PC3 prostate cancer cells treated with the MYC inhibitor MYCi361 were downloaded from NCBI GEO GSE135876 [41]. Published results of ECHO analysis of a circadian time-series of macrophages were downloaded from supplementary material of a publication (doi: 10.1101/gr.263814.120); RNA-sequencing was processed and mapped / aligned as described [9].

### Processing of raw RNA-sequencing data

For U2OS both replicates, raw reads were processed by the University of Pennsylvania Next Generation Sequencing Core to demultiplex using bcl2fastq2-v2.17.1.14. For SHEP both replicates and SKNAS replicate 1, raw reads were processed by Novogene to demultiplex, remove reads containing adaptors, remove reads containing N > 10% (N represents the base cannot be determined), and remove reads containing low quality (Qscore<= 5) base which is over 50% of the total base. For SKNAS replicate 2, raw reads were processed by the GRC to demultiplex, then with FASTP to trim adaptors and remove reads that were low quality or too short. MEF and PC3 RNA sequencing data were processed as previously described [41, 43]. All processed reads were mapped to transcripts using Salmon 1.3.0 [107] in mapping-based mode for paired-end samples, using a decoy-aware transcriptome built from Gencode v35 GRCh38 primary assembly genome and v35 transcriptome. Transcripts were collapsed to gene-level using Tximport 1.12.3 [108] using Gencode v35 transcriptome. Genes were annotated with symbols using the Ensembl GRCh38.101 transcriptome annotations. Tximport yielded as outputs raw counts, which were used for differential expression analysis, and read and length-normalized TPM (Transcripts per million), which were used for ECHO circadian rhythm analysis.

### Metabolite collection and mass-spectrometry

U2OS MYC-ER were plated at 100,000 cells / mL. 24 hours later, cells were treated with ethanol or 4-hydroxytamoxifen to activate MYC-ER or N-MYC-ER. 24 hours later, cells were treated with 0.1 µM dexamethasone (Sigma) to entrain the molecular circadian clock. 24 hours after dexamethasone treatment, cells were gently scraped from the plate in ice-cold PBS (Gibco), pelleted, and snap frozen. Polar metabolites were extracted with methanol and chloroform and redissolved after drying into an acetonitrile:water mixture using a modified Bligh-Dyer method, as previously described [10, 109, 110]. Solubilized polar metabolites were subject to ultraperformance liquid chromatography (UPLC) on a Waters Acquity UPLC coupled to a Waters TQD mass spectrometer (Waters Corporation) as previously described [10, 111].

Following UPLC, mass spectrometry was performed on a QQQ quadrupole instrumentation, (Xevo TQD or TQS-micro, Waters Corporation), as previously described [10, 111]. Specific metabolites were targeted with multiple reaction monitoring, validated against standards and / or mass-spectrometry databases. Note that the panel of standards was heavily altered between when Replicate A and Replicate B were run; hence, these experiments are presented as individual replicates rather than being averaged. Each sample was acquired multiple times, and the sample queue was randomized to remove bias [112]. For each Replicate, the entire set of injections was bracketed by standard metabolites at both the beginning and the end of the run to evaluate instrument performance, sensitivity, and reproducibility.

Mass spectrometry output data were processed as previously described [10], using Waters TargetLynx software (version 4.1), with ion counts processed in R using a custom script. To account for instrument drift, quality control samples, which were a pool of all samples, were injected at the start of every batch for UPLC column equilibration and every 6 injections during mass spectrometry analysis. To normalize each metabolic feature, a LOESS function (locally-weighted scatterplot smoothing) was fitted to the QC data. These values were used for ECHO analysis of circadian rhythmicity. Amino acid quantitation presented in **Figure 7C** and **Supplemental Figure 7** were further normalized to cell counts, which were taken every six hours during the time-series described above. Missing cell counts were interpolated from neighboring values.

### Circadian analysis with ECHO, and graphing of circadian data in the R programming language

The Extended Circadian Harmonic Oscillator (ECHO) application [38] was chosen for oscillation analysis of RNA-sequencing and metabolomics data because it uses parametric approaches to determine rhythmicity in dampening oscillators, which are commonly observed in cell lines. ECHO v4.0 was used. *For qPCR data, RNA-sequencing data, and metabolomics data,* the following parameters were used: paired replicates, unexpressed genes or metabolites removed, linear trends removed, and default parameters for determined if a gene or metabolite is harmonic, overexpressed, or repressed. Genes or metabolites were determined to have circadian rhythmicity if they had a period between 20-28 hours (unless otherwise noted in **Figure 1)**, if the genes were not overexpressed or repressed (ie, sharply increasing or decreasing over the time series by more than an amplitude change coefficient of 0.15), and had a p < 0.05, and a BH.Adj.P < 0.05, except for SKNAS RNA-sequencing. For SKNAS RNA-sequencing, since the two circadian time-series were performed 11 years apart (2012 and 2023) with different SKNAS N-MYC-ER lineages, different RNA isolation techniques, and different RNA-sequencing methods (150 bp vs 100 bp), a slightly more relaxed threshold of a p < 0.05, and a BH.Adj.P < 0.1 was used. *For protein expression data*, the following parameters were used: paired replicates, linear trends removed, smoothing enabled, a tight window of 22-26 hour oscillations, and otherwise default parameters. Proteins were considered oscillatory with a p < 0.05, and a BH.Adj.P < 0.05.

### Graphing of circadian data

For heatmap data in **Figures 2 and 8** and **Supplementary Figures 2 and 8**, data were scaled using the Rescale R package to between -2 and 2 for each gene or metabolite, so they could be compared to each other, and then were sorted by phase (‘hours.shifted’). Gene level data are graphed as heatmaps in ggplot2 using the civvidis colorblind-compliant color scale, and are presented as detrended TPM values (‘original’), while metabolites are presented as computed cosine fit (‘smoothed’), as previously performed [31]. For amplitude comparisons between cell lines and primary cells in **Supplemental Figure 2A,** median amplitude (‘initial.amplitude’) was calculated for all significant oscillating genes, and graphed with 95% confidence intervals in GraphPad Prism 9.0. For amplitude analysis of MYC-OFF vs MYC-ON in **Supplemental Figure 2C**, amplitudes (‘initial.amplitude’) of significant oscillators as defined above were plotted with ggplot2 using a mirrored histogram and logarithmic scale. For dot and line plots in **Figure 8** and **Supplemental Figures 3 and 8**, dots represent detrended TPM values or normalized abundance metabolite values, and lines represent computed cosine fit, as previously performed [31]. Graphs of protein abundance in **Figure 6** represent averages, with error bars representing S.E.M., of detrended and smoothed protein abundance samples that were previously normalized to Tubulin controls.

### Differential expression with DeSeq2

Differential expression of counts data from MYC-ON vs MYC-OFF was performed using DeSeq2 v1.24.0 [42]. For each cell line, every timepoint from each replicate, where applicable, was used as a biological replicate for MYC-OFF and MYC-ON. The DeSeq2 design included time and condition, the alpha cutoff for significant enrichment was set at 0.05, and the apeglm algorithm [113] was used to normalize and shrink the results (‘lfcshrink’). Genes with a padj ≤ 0.05 log2-fold change between MYC-ON vs MYC-OFF were determined to be significant and used for further analysis. Differential expression of MYCi361-treated PC3 cells vs control-treated cells [41] was performed on biological replicates using DeSeq2 v1.24.0 with the same strategy, except that time was not included in the DeSeq2 design.

### Pathway enrichment workflow: Gene Set Enrichment Analysis (GSEA)

Gene Set Enrichment Analysis (GSEA) was performed using GSEA software v4.0 and the Molecular Signatures Database Human Genesets v7 [56-59]. Genes with a log 2 fold change ≥ ± 0.15, representing a 10% or greater change, and a padj ≤ 0.05, were subsetted for GSEA PreRanked, using default parameters. The 10% or greater change cutoff was used because GSEA tests potentially small changes across entire pathways, which may not be represented if only large gene expression changes are included [56-59]. Gene sets with less than 15 genes or more than 500 genes were excluded. GSEA results with an FDR q-value ≤ 0.25 were determined to be significant, as was previously described [56]. Results from molecular signature databases that were not available for Phase Set Enrichment Analysis (below) and for micro RNAs were excluded from analysis for purposes of this manuscript, but full results are available online at FigShare (https://doi.org/10.6084/m9.figshare.c.6754206). GSEA results are presented as normalized enrichment score (NES).

### Pathway enrichment workflow: Phase Set Enrichment Analysis (PSEA)

Phase Set Enrichment Analysis (PSEA) was used to determine which pathways were enriched for genes that oscillated with a similar phase [46]. For each ECHO analysis (MYC-OFF and MYC-ON for SHEP, SKNAS, and U2OS), phases (‘hours.shifted’) from genes determined to have significant circadian oscillation were used along with gene names for PSEA analysis. PSEA v1.1 was run using default parameters, querying for oscillations up to 28 hours, and using the following Molecular Signatures Databases (all v7): C2 (curated gene sets), C3 (regulatory target gene sets), C5 (ontology gene sets), C6 (oncogenic signature gene sets), C7 (immunologic signature gene sets), and H (hallmark gene sets). Gene sets with a Kuiper q-value (vs. background) < 0.2 and p-value (vs. background) <0.1 were determined to be significant. As with GSEA, micro RNA-related results were excluded from analysis for purposes of this manuscript, but full results are available online at FigShare (https://doi.org/10.6084/m9.figshare.c.6754206). For plotting on a polar histogram, results were binned by phase (rounded to the nearest whole number), and plotted on a 28-hour histogram using ggplot2. For bar charts, -logP was computed from the p value of significantly enriched sets, and plotted. For instances where the p value was equal to 0, the –log p value was arbitrarily set to be less than 1 unit higher than the next most significant value in the set.

### Pathway enrichment workflow: ToppFun

ToppFun functional enrichment [49] was used as an alternate method for pathway enrichment of circadian oscillating genes, and genes regulated by MYC. For circadian oscillating genes, all genes that were determined to be significantly rhythmic in each cell line in MYC-OFF or MYC-ON conditions (see “Circadian analysis with ECHO”, above) were entered into ToppFun. For genes regulated by MYC, DeSeq2 for MYC-ON vs MYC-OFF was subsetted for genes with a log 2 fold change ≥ ± 0.66, representing a 1.5-fold or greater change, and a padj ≤ 0.05. In all cases, the default background set for each category was used. When an inputted gene was not found, the default suggested alternative was used. Gene sets were limited to between 5 and 500 genes. An FDR.Q of ≤ 0.05 was considered to be significant. For this manuscript, only GO and Pathway analyses were considered and graphed, but full results are available online at FigShare (https://doi.org/10.6084/m9.figshare.c.6754206). For bar charts, -logP was computed from the p value of significantly enriched sets, and plotted. For instances where the p value was equal to 0, the –log p value was arbitrarily set to be less than 1 unit higher than the next most significant value in the set.

### Pathway enrichment workflow: MetaboAnalyst enrichment analysis

MetaboAnalyst enrichment analysis [114] was used to determine enriched pathways among circadian metabolites. Metabolites determined to be circadian in MYC-OFF or MYC-ON conditions (see “Circadian analysis with ECHO”, above) were manually curated by peak phase, as determined by heatmap analysis, into different groups (CT10 and CT20 for MYC-OFF for both Replicates, CT4 and CT16 for MYC-ON Replicate 1, and CT0, CT8, and CT12 for MYC-ON Replicate 2). For each group, KEGG IDs [115] were inputted and compared against default background. Where multiple KEGG IDs existed for a single mass spectrometry peak, the first KEGG ID was used. Metabolites were compared against the KEGG Metabolite set library. Only results with two or more enriched metabolites were used. p ≤ 0.05 was used to determine significance, and – log p is graphed as bar charts. For plotting on a polar histogram, results were binned by phase, and plotted on a 28-hour histogram using ggplot2.

### Statistical analysis

Statistical analysis and tests for significance for ECHO, GSEA, PSEA, DeSeq2, ToppFun, and MetaboAnalyst Enrichment Analysis are described above. For statistical analysis of total protein expression levels, On-cell western, and mass spectrometry quantitation of amino acids, we used Welch’s corrected Student’s t-test (‘Welch’s t-test’, which does not assume equal variance of sample data), with p ≤ 0.05 determined to be significant and marked with an *.

### Raw data, processed data, and code availability

RNA-sequencing data have been uploaded to NCBI GEO and are available at GEO SuperSeries GSE221174, which contains GSE221103 (SHEP both replicates, SKNAS replicate 1), GSE221173 (U2OS both replicates), and GSE237608 (SKNAS replicate 2). For the purposes of this study, we focused on the “top hits” from each of our analyses (PSEA, GSEA, etc), and present them as ranked lists. However, we identified hundreds to thousands of genes and pathways that oscillate in MYC-OFF cells, and thousands of genes and pathways that were up- or down-regulated by MYC when MYC is activated (MYC-ON). For those researchers who wish to probe a specific gene or pathway of interest that was not listed in our top hits, complete tables and all input data (including metabolomic data) are freely available online at FigShare (https://doi.org/10.6084/m9.figshare.c.6754206).

### Statement on colorblind compliance

All color schemes have been specifically chosen to be easily distinguishable by those with any variety of colorblindness. Colors for graphs and Venn diagrams were chosen using Colorblind Universal Design (https://jfly.uni-koeln.de/color/, acknowledgements to Dr. Masataka Okabe of the Jikei Medial School, Japan, and Dr. Kei Ito of the University of Tokyo, Institute for Molecular and Cellular Biosciences, Japan). Heat maps in R were generated using the Cividis color scale, which is adapted to colorblindness [116].

## Acknowledgements

We would like to acknowledge Hannah De los Santos and Dr. Jennifer Hurley (Rensselaer Polytechnic Institute) for assistance with implementing the ECHO algorithm, Dr. Aimee Edinger (University of California Irvine) for helpful discussion on nutrient transporters, Dr. Lin Zhang (University of Pennsylvania Perelman School of Medicine) for helpful discussion on analysis of RNA-sequencing data, Dr. Fabio Hecht Castro Medeiros (University of Rochester Medical Center) for assistance on design and creation of a graphical abstract, Dr. Adam Wolpaw (Children’s Hospital of Philadelphia) for provision of SKNAS N-MYC-ER cells, and Dr. Zachary Chalmers (Feinberg School of Medicine, Northwestern University) for assistance with downloading publicly available data of PC3 cells treated with MYCi361 [41]. We would also like to acknowledge the University of Rochester Wilmot Cancer Institute Genomics Research Center sequencing services for SKNAS replicate 2, and for guidance on processing and analyzing RNA-sequencing data and implementing differential expression analysis. This work was supported by F30CA200347 and T32CA009140 (to ZEW), F31CA232551 (to RB), R21CA213234 and R01DK120757 (to AMW), R01CA057431 and R01CA051497 (to CVD), and R00CA204593 (to BJA). SSM and BJA were also supported by the Wilmot Cancer Institute.

## Author Contributions

**Conceptualization**: J.C., A.M.W., C.V.D., and B.J.A.; **Methodology**: R.E.D., A.L.H., A.M.W., and B.J.A.; **Software**: R.E.D., S.K., J.B.B., D.M., R.B., S.S.M., A.M.W., and B.J.A.; **Validation**: J.C., R.E.D., S.K., A.M.W., and B.J.A.; **Formal analysis**: J.C., R.E.D., J.B.B., A.L.H., R.B., and B.J.A.; **Investigation**: J.C., R.E.D., S.N.A.B.A.S., J.B.B., D.M., S.K., A.L.H., Z.E.W., R.B., and B.J.A.; **Resources**: A.M.W., C.V.D., and B.J.A.; **Data Curation**: J.C., D.M., S.K., A.M.W., and B.J.A.; **Writing - Original Draft**: J.C. and B.J.A.; **Writing - Review & Editing**: J.C., J.B.B., Z.E.W., R.B., S.S.M., A.M.W., C.V.D., and B.J.A.; **Visualization**: J.C., R.E.D., J.B.B., S.K., R.B., and B.J.A.; **Supervision**: A.M.W., C.V.D., and B.J.A.; **Project administration**: R.E.D., A.M.W., C.V.D., and B.J.A.; **Funding acquisition**: Z.E.W., R.B., A.M.W., C.V.D., and B.J.A.

## Supplemental Figures and Tables

**Supplemental Figure 1. MYC and N-MYC are overexpressed in MYC-ER and N-MYC-ER-expressing cells, and naturally overexpressed MYC controls circadian gene expression.** A. *MYCN* was determined in parental SKNAS, SKNAS N-MYC-ER, or SHEP N-MYC-ER by qPCR in n=3 replicates by quantitative PCR (qPCR), normalized to β2M. *MYCN* overexpression is shown relative to parental SKNAS, since SHEP do not express *MYCN.* B. C-MYC-ER or endogenous c-MYC (endo) are shown in n=3 immunoblot replicates, and quantitation is shown in the right panel, relative to Tubulin. **C.** The PC3 prostate cancer cell line, known to express high MYC, was treated with 6 µM of the MYC inhibitor MYCi361 for 24 hours in biological triplicates, as previously published [41]. Raw RNA-sequencing data was downloaded and processed (see **Methods**), and Deseq2 was used to compare cells ± MYCi361. Log2FC of indicated genes is shown. For **A,B**, error bars are standard error of the mean (S.E.M.) and ** is p < 0.01 and * is p < 0.05 by Welch’s Corrected Student’s T-test. For **C**, error bars are Standard Error as calculated by DeSeq2, and * indicates padj < 0.005.

**Supplemental Figure 2. Comparison of MYC-OFF oscillation to transcriptomic oscillation in primary cells, and impact of MYC on gain of oscillation. A.** The median initial amplitude (absolute, abs) of oscillating genes in SHEP, SKNAS, and U2OS MYC-OFF, as determined by ECHO and shown in **Figure 2**, was compared to previously published ECHO analysis of time-series RNA-sequencing of entrained primary macrophages (n=3 biological replicates) or new ECHO analysis of previously published and downloaded time-series RNA-sequencing of entrained primary mouse embryonic fibroblasts (n=4 biological replicates, see **Methods**) [9, 43]. Error bars represent 95% confidence intervals. **B.** RNA-sequencing was performed on SHEP N-MYC-ER, SKNAS-N-MYC-ER or U2OS MYC-ER ± 4OHT and + dexamethasone, with RNA samples collected every 2-4 hours at the indicated timepoints. RNA was analyzed for rhythmicity by ECHO for both MYC-OFF and MYC-ON, with genes with a 20-28 hour period and with BH.Adj.P.Value < 0.05 deemed to be rhythmic. These genes were sorted by phase and are presented in a heatmap for MYC-ON. For MYC-OFF, the same genes that are rhythmic in MYC-OFF are presented in the same order, but with MYC-OFF values instead. N=2 time series were used for each cell line. **C.** The amplitude of oscillation of each gene in MYC-OFF and MYC-ON, as determined by ECHO, was graphed as a mirrored density plot to allow direct comparison between each condition, with MYC-OFF on the top and MYC-ON on the bottom.

**Supplemental Figure 3. MYC induces an alternate oscillatory program that includes cell cycle. A.** The pathways deemed to be significantly oscillatory in MYC-ON cells by PSEA (20-28 hr period, q-value (vs. background) < 0.2 and p-value (vs. background) <0.1) for each cell line were binned and graphed on a polar histogram. The scale for each histogram is on the left side. Overlap of oscillatory programs by Venn diagram is also shown. **B.** The identity of oscillatory programs in MYC-ON cells from (**A**) are plotted by average of –LogP from each cell line of overlap. The most highly significant pathways (up to 10) for each overlap are shown. **C,D.** Cell cycle genes deemed to be oscillating by ECHO for MYC-OFF and MYC-ON SHEP (**C**) and U2OS (**D**) are graphed, with dots representing the RNA-sequencing TPM values, and lines indicating the fitted oscillation curves as calculated by ECHO. Genes with a green border are oscillatory in both MYC-OFF and MYC-ON.

**Table 1. An independent methodology reveals oscillating gene pathways in MYC-inducible cells.** Genes from SHEP N-MYC-ER, SKNAS-N-MYC-ER or U2OS MYC-ER MYC-OFF or MYC-ON that were deemed oscillatory by ECHO, without regards to their phase, were analyzed for pathway enrichment with the ToppFun suite, and pathways with FDR B&H < 0.05 were deemed significant. The most highly enriched pathways from the Pathway and GO libraries are shown in the table, ranked by –LogP.

**Supplemental Figure 4. The most altered genes by MYC correlate with changes in biosynthesis, metabolism, and cell attachment. A.** Differential expression analysis, using DeSeq2, was performed on MYC-ON vs MYC-OFF for SHEP N-MYC-ER, SKNAS-N-MYC-ER or U2OS MYC-ER, with p.adj < 0.05 deemed significant. Genes that were up- or down-regulated in MYC-ON by at least 1.5-fold are shown. **B,C.** Venn diagram of the overlap of 1.5-fold upregulated (**B**) and downregulated (**C**) genes by MYC. **D, E.** Genes that were upregulated or downregulated in all 3 cell lines were subjected to pathway analysis with the ToppFun suite, and pathways with FDR B&H < 0.05 were deemed significant. For upregulated (**D**) and downregulated (**E**) genes, the most highly enriched pathways from the Pathway and GO libraries are shown in the table, ranked by –LogP.

**Supplemental Figure 5. A gene-centric methodology reveals pathways that shift from oscillatory to MYC-regulated. A.** Illustration of workflow to identify genes that lose oscillation when MYC is activated (by ECHO), which of these genes become up- or down-regulated by MYC (using DeSeq2), and analysis of these genes for pathway enrichment by ToppFun. **B-D.** Venn diagram from SHEP N-N-MYC-ER (**B**), SKNAS N-MYC-ER (**C**), or U2OS MYC-ER (**D**) of genes (identified by ECHO) that were circadian in MYC-OFF (center, green), and lost oscillation and became either downregulated (left, pink) or upregulated (right, tan) in MYC-ON (identified by DeSeq2). Bottom section shows the most highly enriched upregulated pathways in each cell line, as identified by ToppFun enrichment from the Pathway and GO libraries of genes that lost oscillation and became upregulated. Pathways are ranked by –LogP, and FDR B&H < 0.05 for all pathways. For SKNAS, the most enriched pathways that correspond to the molecular circadian clock are not shown, to instead focus on output pathways.

**Supplemental Figure 6. MYC upregulates protein levels of several amino acid transporters, and affects molecular clock protein levels**. Immunoblots in **Figure 6** were quantified, relative to Tubulin, for each indicated protein, and MYC-ON was compared to MYC-OFF for each band and each replicate regardless of time. Error bars are S.E.M, and **** is p < 0.0001, *** is p < 0.001, and * is p < 0.05 by Welch’s corrected Student’s T test.

**Supplemental Figure 7. MYC enhances intracellular amino acid pools.** A replicate experiment (Replicate B) of **Figure 7C** was performed. LC-Mass spectrometry was performed on U2OS MYC-ER treated ± 4OHT for at least 48 hours. N=25 circadian timepoints for MYC-OFF and MYC-ON were averaged as biological replicates, normalized to cell number for each collection. **** is p < 0.0001, *** is p < 0.001, ** is p < 0.01, and * is p < 0.05 by Welch’s corrected Student’s T test.

**Supplemental Figure 8. MYC disrupts cell autonomous oscillations in a replicate experiment, but induces alternate oscillations. A.** An independent replicate (‘Replicate B’) of time-series metabolite collection from U2OS MYC-ER cells was performed in an identical fashion to that described in **Figure 6A**. Rhythmicity was assessed by ECHO for both MYC-OFF and MYC-ON, with metabolites with a 20-28 hour period and with BH.Adj.P.Value < 0.05 deemed rhythmic. These metabolites were sorted by phase and are presented in a heatmap for MYC-OFF. For MYC-ON, the same metabolites that are rhythmic in MYC-OFF are presented in the same order, but with MYC-ON values instead. **B.** Metabolites from Replicate B deemed to be oscillating by ECHO for MYC-OFF are graphed, with dots representing the relative abundance values, and lines indicating the fitted oscillation curves as calculated by ECHO. Metabolites with a green border are oscillatory in both MYC-OFF and MYC-ON. **C, D. For Replicate A** (**C**) from Figure 6, and **Replicate B** (**D**), metabolites deemed oscillatory from MYC-ON are shown, sorted by phase. For MYC-OFF, the same metabolites that are rhythmic in MYC-ON are presented, but with MYC-OFF values instead.

**Supplemental Figure 9. MYC disrupts metabolic circadian programming. A,B.** KEGG enrichment analysis was performed on metabolites that peaked in the indicated phases from Replicate B in MYC-OFF (**A**) or MYC-ON (**B**) conditions, and significantly enriched pathways were graphed on a polar histogram. For each histogram, the scale is on the left side. Pathways with a p < 0.05 were deemed significant and are graphed.

