## Supplemental Figures and Tables for "MYC disrupts transcriptional and metabolic circadian oscillations in cancer and promotes enhanced biosynthesis"

Supplemental Figure 1

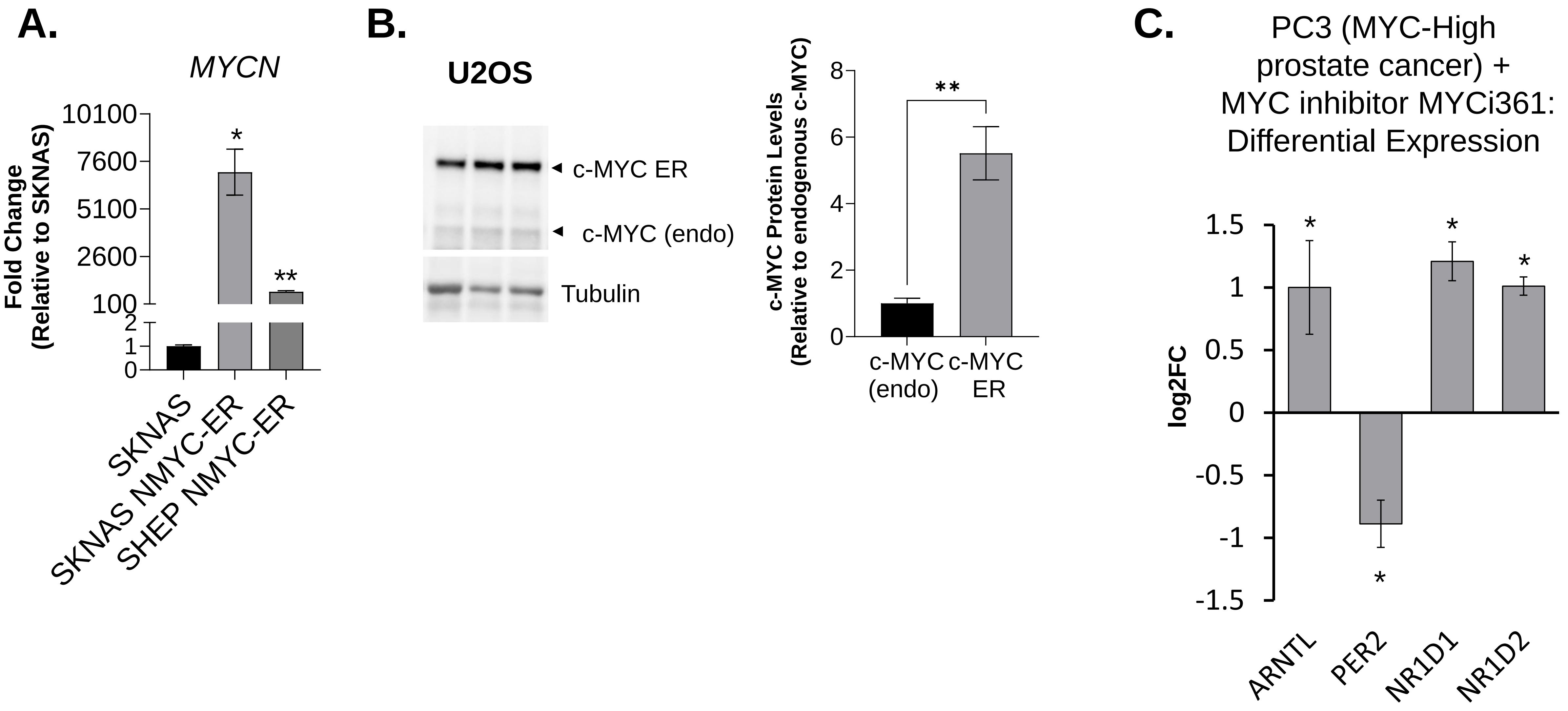

### Supplemental Figure 2

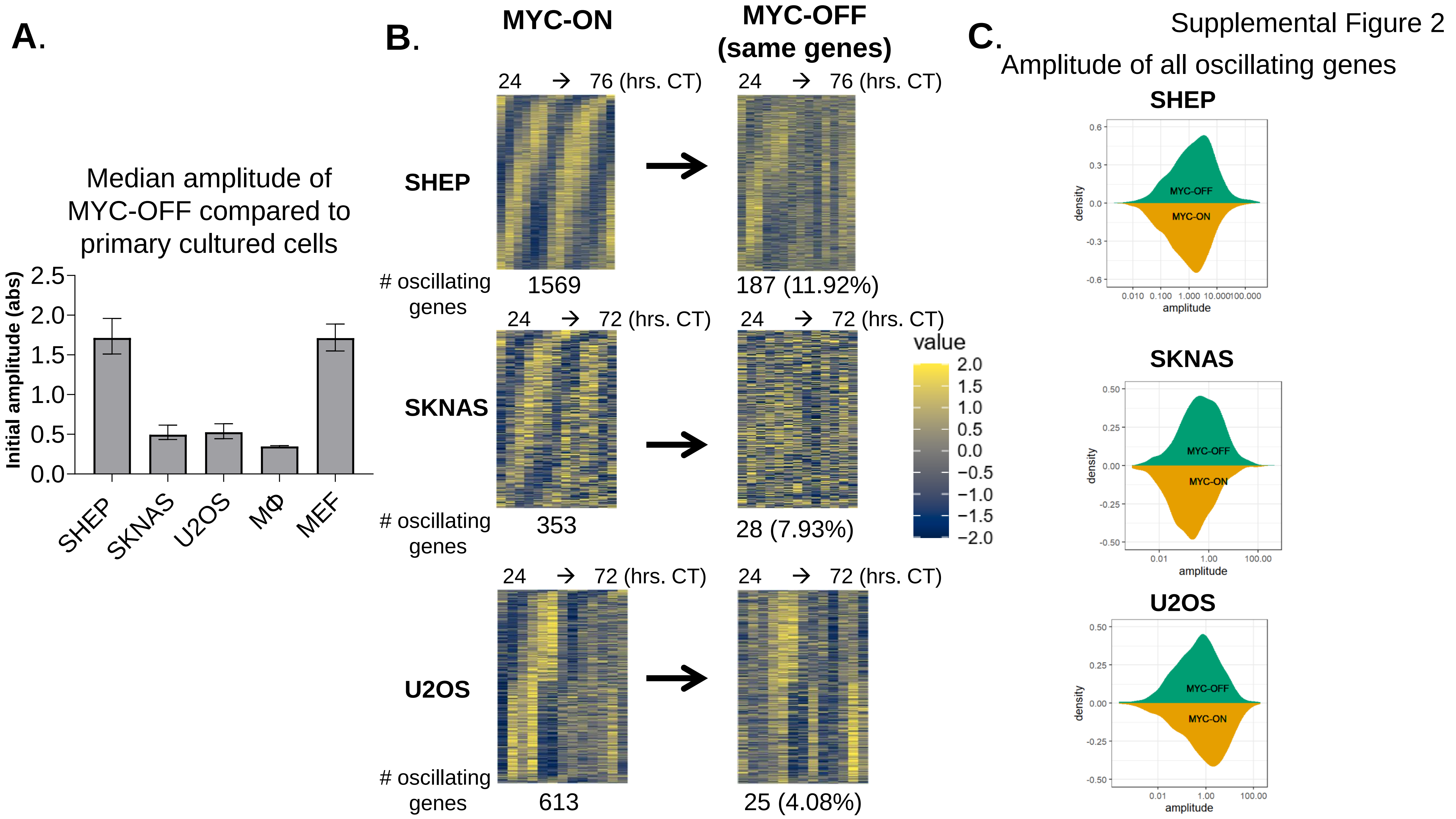

Supplemental Figure 3

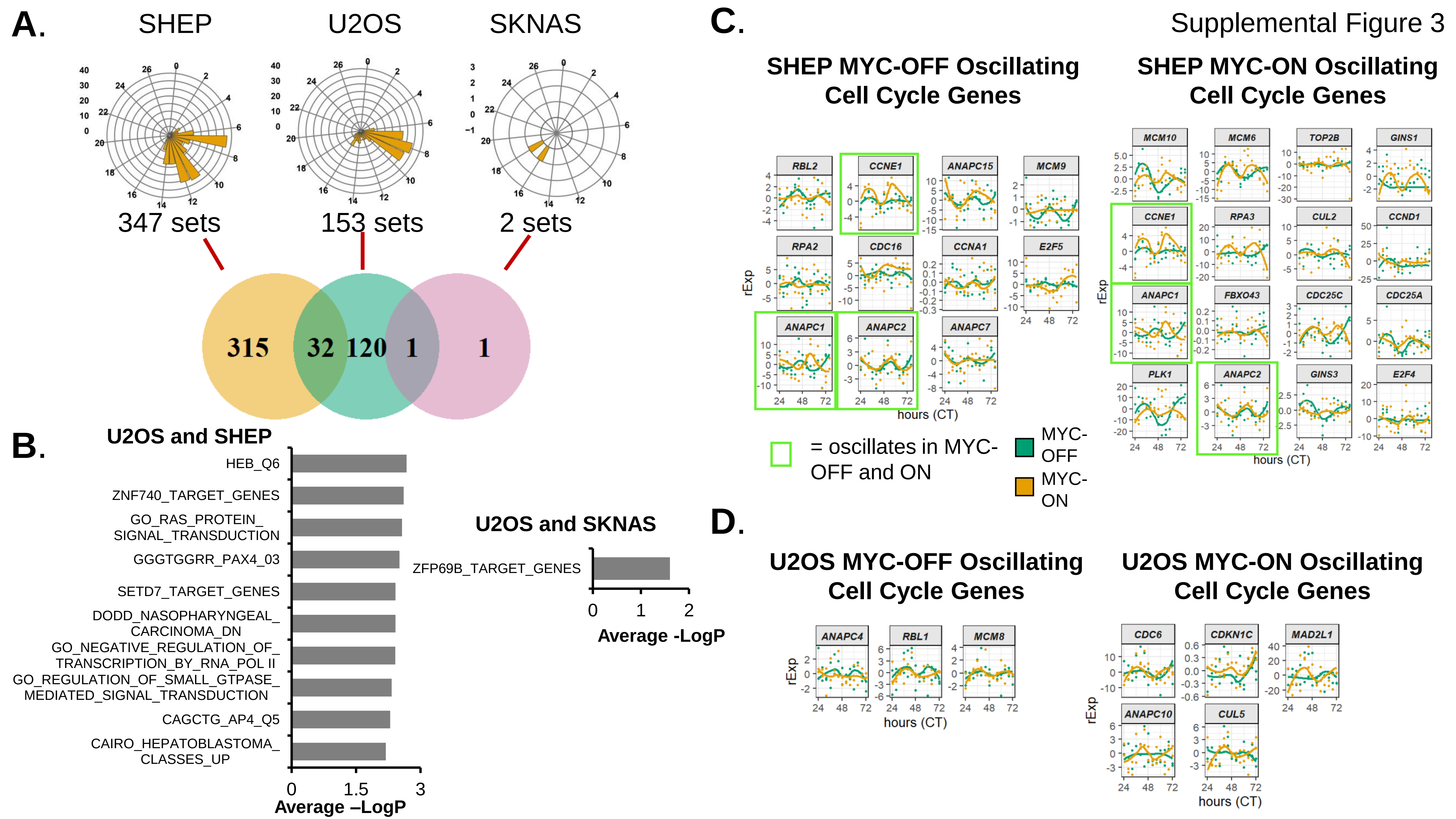

Table 1

| SHEP MYC-OFF | | | | |
|---|---|---|---|---|
| Rank | Category | Name | P value | -log10 P |
| 1 | Pathway: Pathway Interaction Database (PID) | Class I PI3K signaling events | 7.26E-07 | 6.14 |
| 2 | GO: Cellular Component | lysosomal membrane | 4.74E-06 | 5.32 |
| 3 | GO: Cellular Component | lytic vacuole membrane | 4.74E-06 | 5.32 |
| 4 | GO: Cellular Component | vacuolar membrane | 8.01E-06 | 5.10 |
| 5 | GO: Cellular Component | late endosome | 4.33E-05 | 4.36 |
| 6 | GO: Cellular Component | late endosome membrane | 1.43E-04 | 3.85 |
| **SHEP MYC-ON** | | | | |
| **Rank** | **Category** | **Name** | **P value** | **-log10 P** |
| 1 | GO: Biological Process | mitotic metaphase plate congression | 9.16E-06 | 5.04 |
| 2 | GO: Biological Process | metaphase plate congression | 9.46E-06 | 5.02 |
| **SHEP MYC-OFF** | | | | |
| **Rank** | **Category** | **Name** | **P value** | **-log10 P** |
| 1 | Pathway: KEGG | Circadian rhythm | 1.26E-10 | 9.90 |
| 2 | Pathway: MSigDB BIOCARTA | Circadian rhythm - mammal | 1.80E-10 | 9.75 |
| 3 | Pathway: REACTOME | BMAL1:CLOCK,NPAS2 activates circadian gene expression | 1.76E-09 | 8.75 |
| 4 | Pathway: PantherDB | Circadian clock system | 1.99E-08 | 7.70 |
| 5 | Pathway: MSigDB BIOCARTA | Circadian rhythm pathway | 5.90E-08 | 7.23 |
| 6 | GO: Biological Process | circadian regulation of gene expression | 1.18E-07 | 6.93 |
| 7 | Pathway: REACTOME | Circadian Clock | 1.92E-07 | 6.72 |
| 8 | GO: Biological Process | rhythmic process | 3.65E-07 | 6.44 |
| 9 | GO: Biological Process | circadian rhythm | 5.63E-07 | 6.25 |
| 10 | GO: Biological Process | regulation of circadian rhythm | 1.14E-06 | 5.94 |
| **U2OS MYC-ON** | | | | |
| **Rank** | **Category** | **Name** | **P value** | **-log10 P** |
| 1 | GO: Biological Process | negative regulation of centrosome cycle | 3.89E-06 | 5.41 |
| 2 | GO: Biological Process | negative regulation of centrosome duplication | 3.89E-06 | 5.41 |
| 3 | GO: Biological Process | regulation of small GTPase mediated signal transduction | 2.51E-05 | 4.60 |
| 4 | GO: Biological Process | entrainment of circadian clock by photoperiod | 4.67E-05 | 4.33 |
| **SKNAS MYC-OFF** | | | | |
| **Rank** | **Category** | **Name** | **P value** | **-log10 P** |
| 1 | GO: Biological Process | entrainment of circadian clock by photoperiod | 2.04E-07 | 6.69 |
| 2 | Pathway: KEGG | Circadian Rhythm - mammal | 3.83E-07 | 6.42 |
| 3 | GO: Biological Process | entrainment of circadian clock | 1.12E-06 | 5.95 |
| 4 | GO: Biological Process | photoperiodism | 1.12E-06 | 5.95 |
| **SKNAS MYC-ON (none)** | | | | |

Supplemental Figure 4

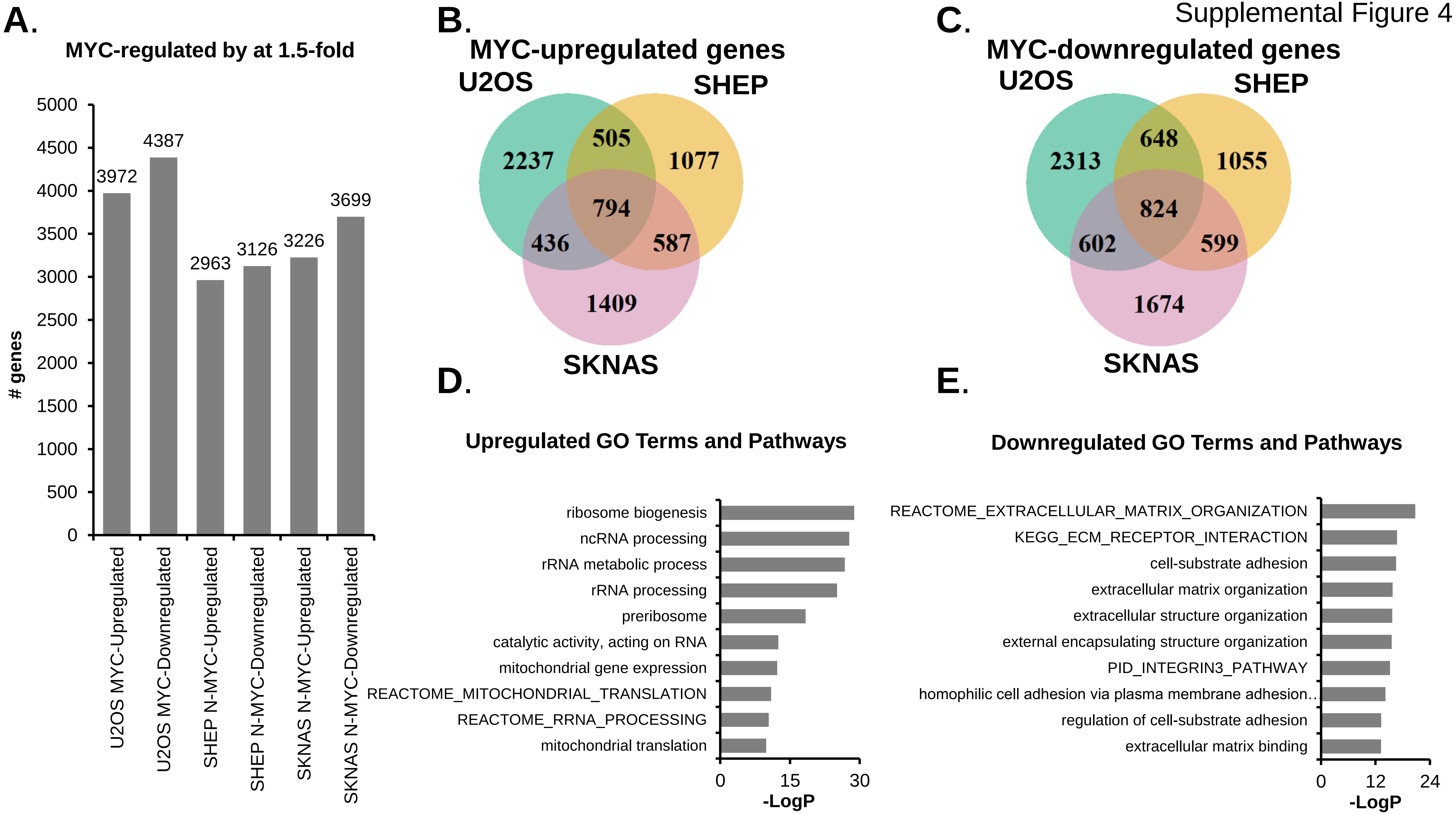

Supplemental Figure 5

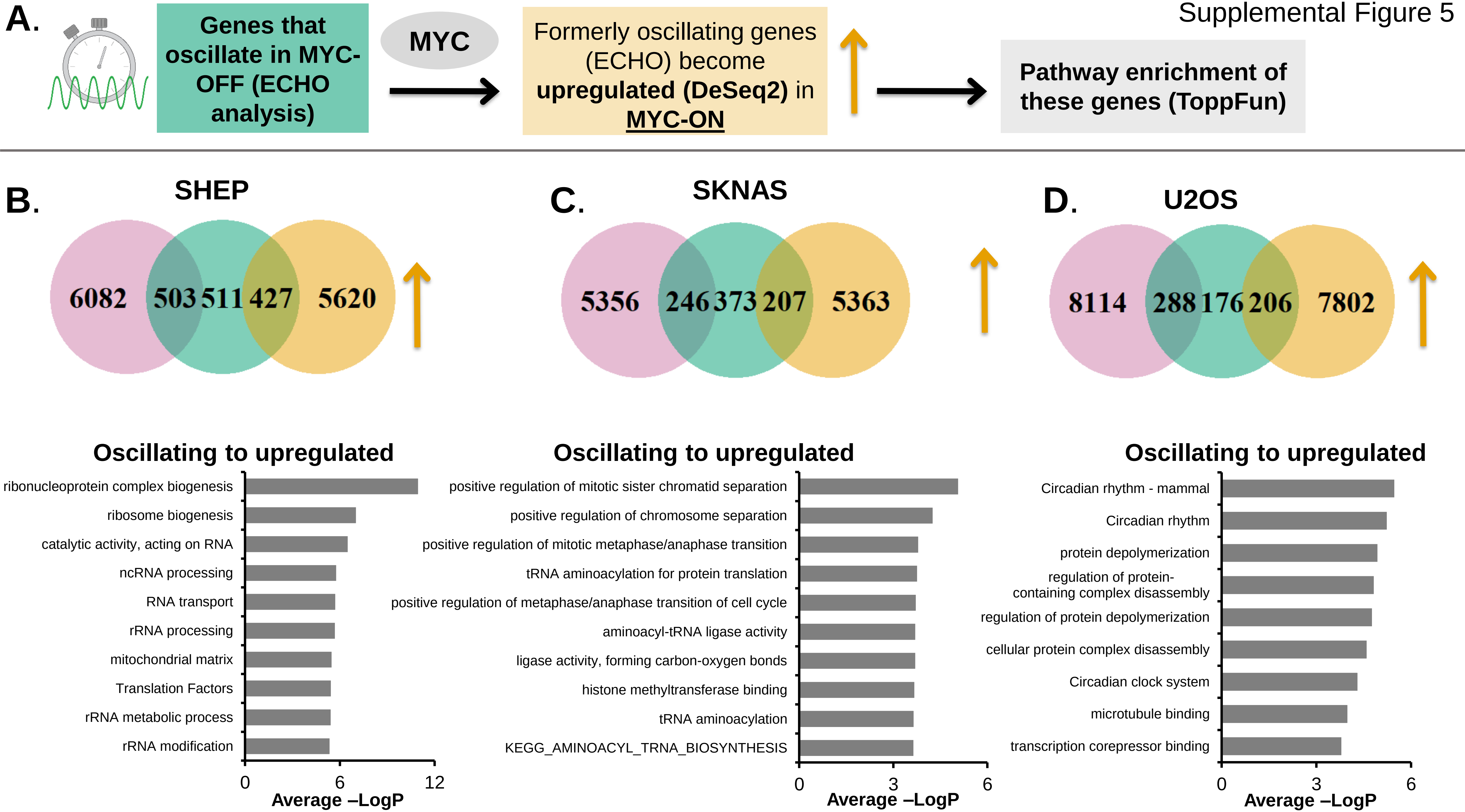

Supplemental Figure 6

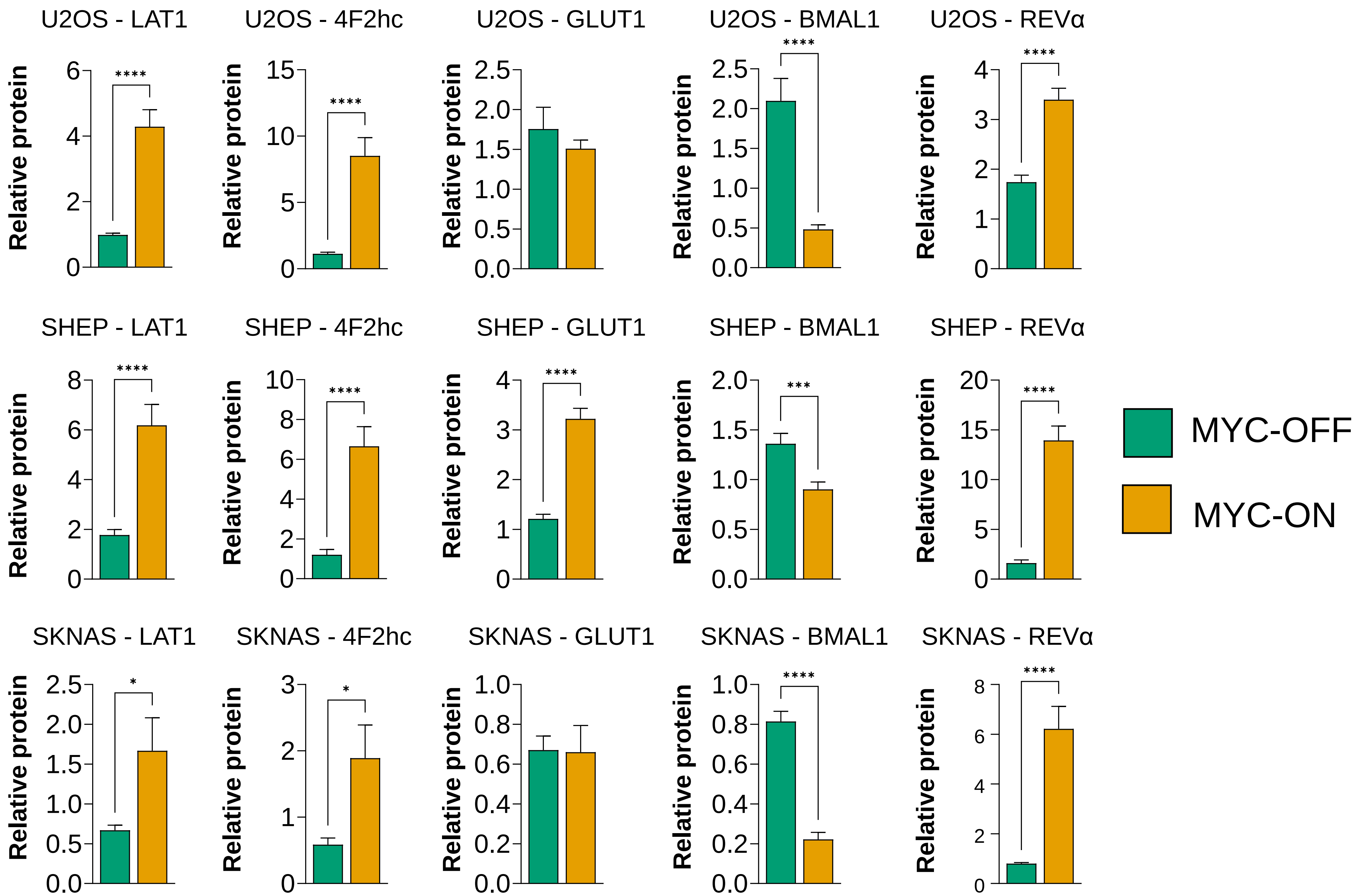

Supplemental Figure 7

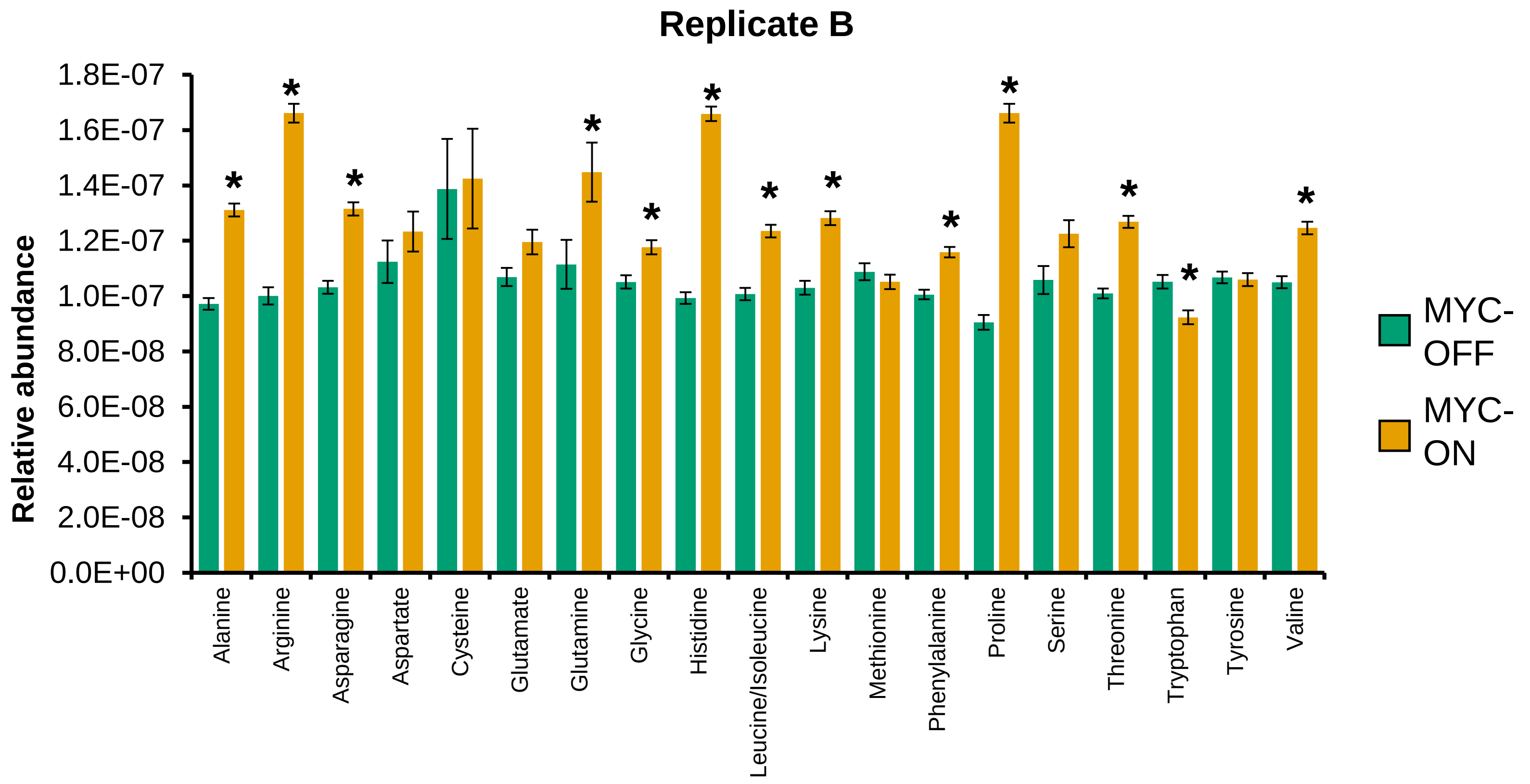

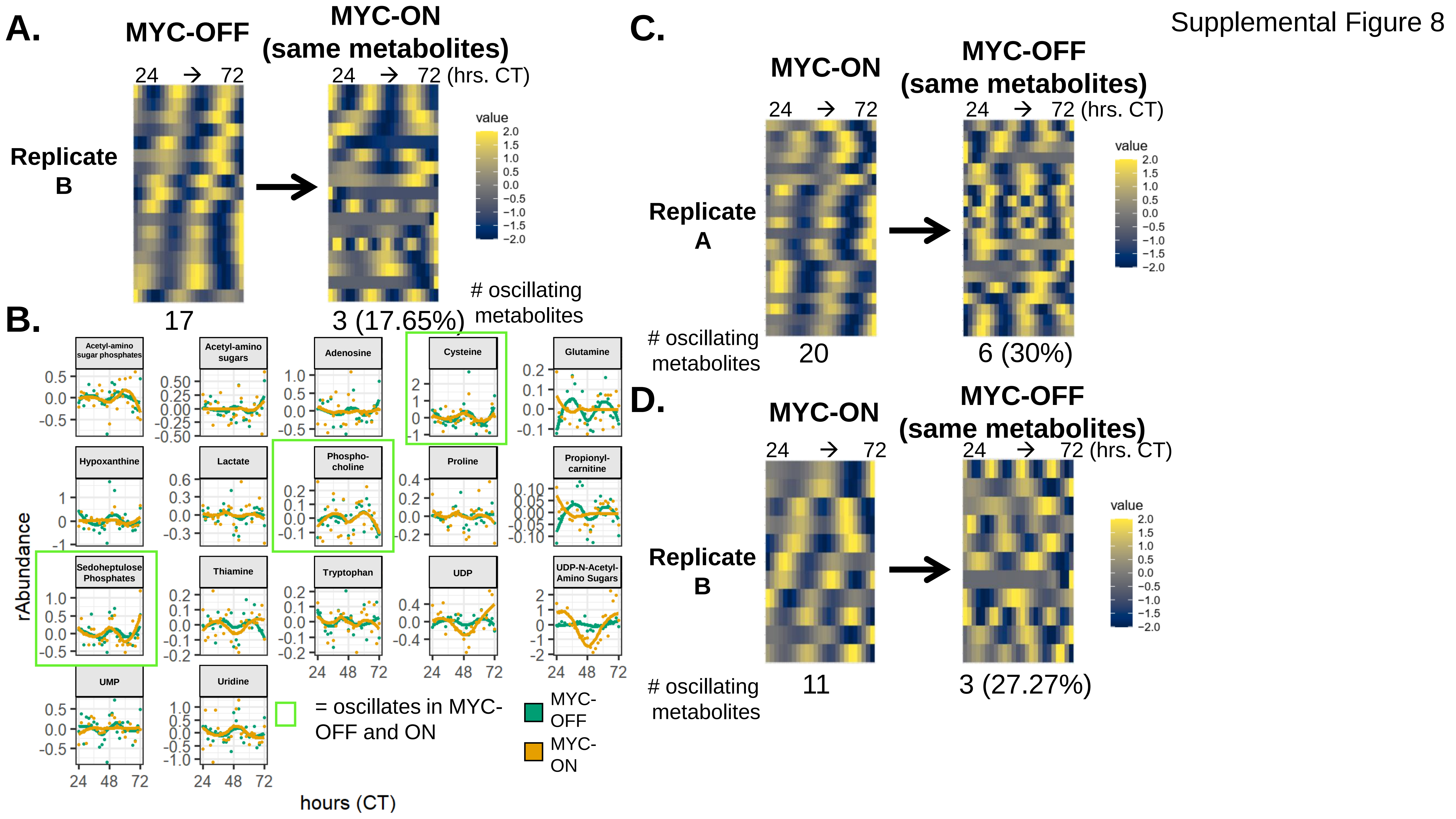

A.
MYC-OFF
MYC-ON (same metabolites)
24 → 72
24 → 72 (hrs. CT)
Replicate B
value
### oscillating metabolites
17
3 (17.65%)
B.
Acetyl-amino sugar phosphates
Acetyl-amino sugars
Adenosine
Cysteine
Glutamine
Hypoxanthine
Lactate
Phospho-choline
Proline
Propionyl-carnitine
Sedoheptulose Phosphates
Thiamine
Tryptophan
UDP
UDP-N-Acetyl-Amino Sugars
UMP
Uridine
rAbundance
hours (CT)
= oscillates in MYC-OFF and ON
MYC-OFF
MYC-ON
C.
MYC-ON
MYC-OFF (same metabolites)
Replicate A
### oscillating metabolites
20
6 (30%)
D.
MYC-ON
MYC-OFF (same metabolites)
Replicate B
### oscillating metabolites
11
3 (27.27%)

### Replicate B: Oscillating Metabolites and KEGG Enrichment of Oscillating Metabolites P<0.05

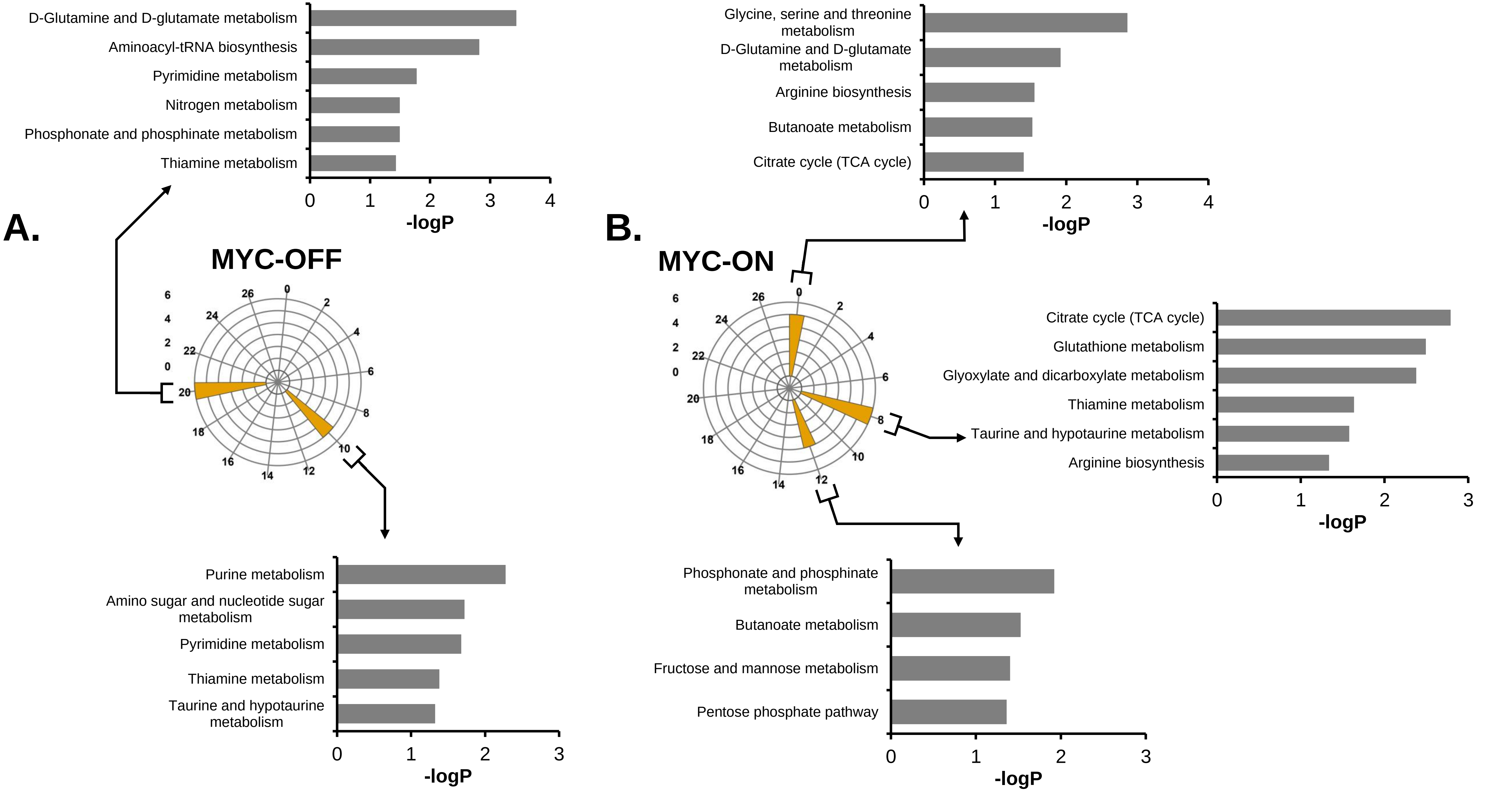
